# Phosphatidylserine and RhoB connect phosphatidylinositol 4-phosphate and phosphatidic acid metabolism at the plasma membrane

**DOI:** 10.1101/2025.09.30.679611

**Authors:** Shiying Huang, Yeun Ju Kim, Xiaofu Cao, Timothy W. Bumpus, Mira Sohn, Matthew Tyler Menold, Ryan S. Dale, Jin Joo Kang, Saori Uematsu, Shagun Gupta, Shu-Bing Qian, Haiyuan Yu, Tamas Balla, Jeremy M. Baskin

**Affiliations:** Department of Chemistry and Chemical Biology, Cornell University, Ithaca, New York, United States, 14853; Weill Institute for Cell and Molecular Biology, Cornell University, Ithaca, New York, United States, 14853; Section on Molecular Signal Transduction, NICHD, National Institutes of Health, Bethesda, Maryland, United States, 20892; Bioinformatics and Scientific Programming Core, NICHD, National Institutes of Health, Bethesda, Maryland, United States, 20817; Department of Computational Biology, Cornell University, Ithaca, New York, United States, 14853; Department of Nutritional Sciences, Cornell University, Ithaca, New York, United States, 14853

## Abstract

Cells tightly control the homeostatic levels and subcellular localizations of membrane phospholipids through the regulation of the activities of numerous lipid-metabolizing enzymes and lipid transfer proteins. Yet, the mechanisms by which lipid imbalances are sensed and corrected to establish and maintain homeostasis are, in most cases, unknown. Here we present an expanded view of plasma membrane (PM) phosphoinositide metabolism by revealing an unexpected metabolic connection between two key anionic lipids in this membrane, phosphatidylinositol 4-phosphate (PI4P) and phosphatidic acid (PA). PM pools of PI4P are generated by PI 4-kinase Type IIIα (PI4KIIIα/PI4KA), an essential enzyme whose partial dysfunction leads to numerous hereditary human diseases. We find that depletion of PI4P by pharmacological inhibition of PI4KA increases the activity of phospholipase Ds (PLDs) and the levels of their lipid product, PA, in the PM. Guided by RNA-seq analysis and proximity labeling proteomics, we elucidate how cells connect this PI4P decrease to a compensatory increase in PA levels. Loss of PM PI4P induces a concomitant decrease of phosphatidylserine (PS) levels, and this metabolic rewiring activates a reciprocal relationship between PS synthesis and PLD-mediated PA generation. These metabolic changes also lead to transcriptional and translational upregulation of the small GTPase RhoB, which enhances PLD-mediated PA synthesis and subsequent actin cytoskeletal remodeling. Our study reveals how disease-relevant perturbation of phosphoinositide synthesis induces an integrated response that ultimately boosts levels of PA, a key anionic lipid and metabolic intermediate in phosphoinositide resynthesis.

## INTRODUCTION

The phosphatidylinositol (PI) cycle, first described by Lowell and Mabel Hokin more than 70 years ago, encompasses the accelerated hydrolysis and resynthesis of phosphoinositides (PPIns) at the plasma membrane (PM) in cells stimulated by segretagogues^1–3^. Subsequent studies found that phosphatidylinositol 4-phosphate (PI4P) and phosphatidylinositol 4,5-bisphosphate (PI(4,5)P₂), the major PPIns in the PM, mediate a plethora of essential cellular functions, including exo/endocytosis, ion channel regulation, calcium signaling, actin organization, and interorganelle lipid transport^4–12^. Further, the increased metabolic flux through the PI cycle in stimulated cells, which generates important lipid and soluble intermediates including diacylglycerol (DAG), phosphatidic acid (PA), PI, and inositol trisphosphate (IP_3_), is a critical component of the signal transduction machinery to alter a variety of cellular functions^13^. Though the PI cycle was originally described in excitable, secretory tissues featuring high rates of exocytosis and compensatory endocytosis, it is a universal feature of all eukaryotic cell types, though rates of flux through the PI cycle can vary considerably depending on cell type and physiological state, e.g., activation of GPCR–Gq or receptor tyrosine kinase signaling, which lead to phospholipase C (PLC)-mediated PI(4,5)P_2_ hydrolysis^14–16^.

An underappreciated feature of the PI cycle is that it is not a closed loop. PI(4,5)P_2_ can be siphoned off into PI(3,4,5)P_3_ via Class I PI 3-kinases, and most prominently, the glycerolipid intermediates DAG and PA are important lipid biosynthetic precursors for triglycerides and all major phospholipid classes that exist outside the core PI cycle. These routes include phosphatidylcholine (PC) and phosphatidylethanolamine (PE) synthesis via the Kennedy pathway (and subsequent conversions to phosphatidylserine (PS)) and production of phosphatidylglycerol (PG) and cardiolipin (CL), in addition to PI, via the CDP-DAG pathway^17^. Because of this built-in leakiness, the core PI cycle (i.e., PI → PI4P → PI(4,5)P_2_ → DAG →PA →CDP-DAG →PI) has the potential to tune the broader lipid biosynthetic network to facilitate optimal adaptation of cellular metabolism during periods of prolonged stimulation. This adaptive response must require additional inputs to maintain diverse phosphoinositide-dependent signaling outcomes while using PI cycle components for building up other lipid components. Yet, despite the long-established nature of the PI cycle, it remains unknown how cells reprogram their lipidome to redirect lipid intermediates to replenish the PI pool in response to changing demands to support adaptive changes and/or during pathological perturbations as occur in various disease states.

Production of PI4P, the immediate precursor to PI(4,5)P_2_, has emerged as a key regulatory point in the PM PPIns metabolic network. Though PI4P is present on several organelle membranes, the PI4P pool at the PM is synthesized primarily by one of four PI 4-kinase isoforms, PI 4-kinase Type IIIα (PI4KIIIα, encoded by PI4KA)^18–20^. Indeed, PI4KIIIα activity is required for acute PI(4,5)P_2_ signaling in cells due to its critical role in PI(4,5)P_2_ resynthesis following its hydrolysis, e.g., by GPCR-Gq-PLC signaling^21,22^. Beyond its roles in acute PLC-mediated signaling, PI4KIIIα and the PM PI4P pool under its control plays crucial housekeeping roles that affect the lipid landscape and regulates development. The PI4P gradient formed between the PM and the ER provides the driving force for the transport of PS from the ER to the PM mediated by members of the oxysterol binding protein related protein family^7,11,23^. This PI4P–PI cycling between the ER and the PM represents a short loop in the context of the original PI cycle, which is evolutionarily more ancient than the receptor-controlled PLC activation pathway.

Accordingly, PI4KIIIα is essential in all tested model organisms. In *S. cerevisiae*, the PI4KIIIα ortholog Stt4 is necessary for cell wall integrity and actin organization^18,24^. In *Drosophila*, PI4KIIIα is required for Hippo signaling and egg chamber polarity and morphology^25,26^. In mice, knockout or chronic inhibition of PI4KIIIα leads to mucosal epithelial degeneration of the gastrointestinal tract, resulting in lethality^21,27^. Mutations in either PI4KA or genes encoding essential non-catalytic components of a PI4KIIIα-containing complex^28–32^ required for PI4P synthesis (EFR3A/B, TTC7A/B, and FAM126A/B) have been linked to schizophrenia^33,34^, hypomyelinating leukodystrophy^31,35,36^, inflammatory bowel disease^37^, and immunodeficiency^36^. Indeed, the importance of PI4KIIIα in cell physiology is further underscored by its essential role during hepatitis C viral infection and replication, where the enzyme is hijacked to produce PI4P on viral replication organelles, consequently antagonizing its vital housekeeping roles at the PM^38,39,27,40,41^.

Collectively, the numerous knockout studies in model organisms and human disease genetics point to chronic deficiency of PM PI4P as leading to catastrophic outcomes at the cellular and organismal levels. However, the downstream events that cause these end results, and the compensatory changes that are initiated under conditions of perturbation or stress, remain largely unknown. Pharmacological inhibition of PI4KIIIα would enable the identification of first-order mechanisms that precede the pleiotropic, long-term effects of PI4KA defects. For example, we recently exploited PI4KIIIα inhibition to reveal key mechanistic insights into how PI4P-dependent interorganelle transport of PS contributes to the complex pathology of Lenz-Majewski syndrome, which is caused by activating mutations in the PS-synthesizing enzyme PSS1^42^.

In this study, we undertook an investigation of the cellular response to disruption of PM PPIns metabolism by pharmacological inhibition of PI4KIIIα. We uncovered an integrated response to PM PI4P depletion that leads to an unexpected accumulation of PA in the PM in large part via activation of phospholipase D (PLD) enzymes, which produce PA via PC hydrolysis. Mechanistically, this response involves a metabolic rewiring wherein PI4P loss causes a loss of PS at the PM, which we find exhibits a reciprocal relationship with PA under conditions of limiting PI4P. These metabolic changes lead to transcriptional upregulation of the small GTPase RhoB, which under PI4P-limiting conditions enhances PLD-mediated PA synthesis and subsequent changes in F-actin organization, explaining perturbations to the actin cytoskeleton observed both by PI4KIIIα inhibition and in genetic models of PI4KA-associated disease. Overall, our findings define a long-range metabolic circuit connecting PI4KIIIα and PLD signaling, revealing unexpected metabolic inputs into PM PPIns metabolism.

## RESULTS

### Inhibition of PI4P synthesis at the PM elevates PA levels

To assess the effect of prolonged PM PI4P depletion on cellular lipid compositions, we treated HEK293 cells overnight with the selective PI4KIIIα inhibitor GSK-A1^21^. Total cellular lipidomics revealed an increase in total PA species upon treatment (Figure 1B). To gain more precise information about the subcellular localization of the PA increase, we used live-cell imaging with the Nir2-LNS2 domain (residues 816-1181) to detect PA and found that it showed prominent increases in the PM following GSK-A1 treatment (Figure 1C). We then used a bioluminescence resonance energy transfer (BRET) assay, which allows for monitoring of organelle-specific lipid changes, to assess PA changes in the PM at the cell population scale. This assay involves expression of a Renilla luciferase variant fused to a lipid-binding domain and an organelle-anchored mVenus construct. The presence of the target lipid on the organelle of interest recruits the luciferase-fused lipid sensor, and subsequent BRET leads to an increase in mVenus fluorescence that serves as a readout for lipid levels on the organelle of interest^43^. We again used the Nir2-LNS2 domain (residues 816-1181) to measure PA at the PM^44^ and found that PI4KIIIα inhibition induced an increase in PA levels at the PM (Figure 1D). Whereas in the classic PI cycle, during PLC activation a DAG kinase (DGK) is downstream of PI4P and converts DAG to PA at the PM (Figure 1A), it is important to note that during these experiments no stimulation of PLC activation was observed, and PI(4,5)P_2_ levels increased rather than decreased upon prolonged GSK-A1 treatment (Figure S2A). Therefore, it was unlikely that PI(4,5)P_2_ hydrolytic product was the source of the elevated level of PA in the PM.

**Figure 1.**
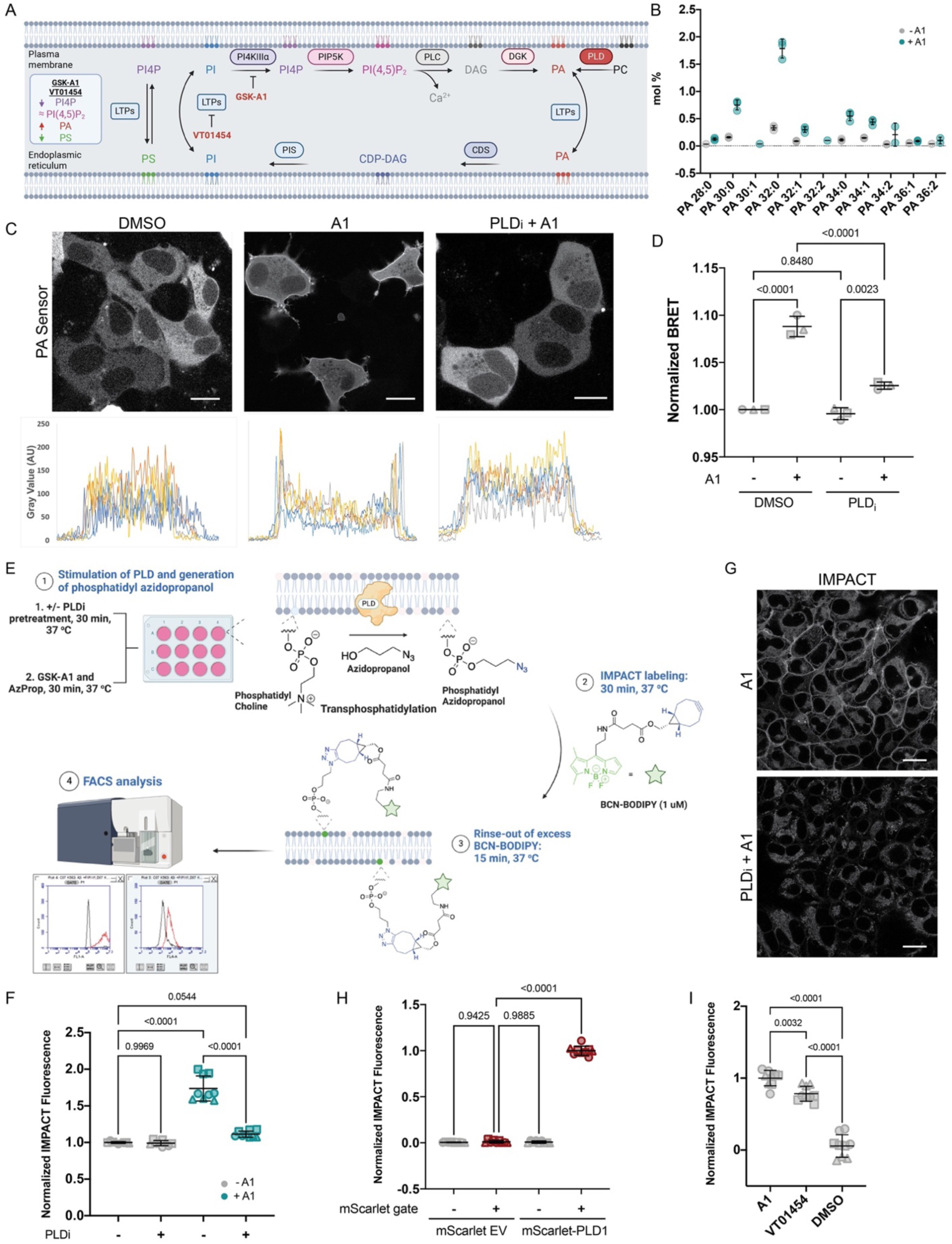
Depletion of PI4P at the PM leads to changes in PA. (A) Schematic depicting the PI cycle and PLD-mediated synthesis of PA. (B) Lipidomics analysis of PA levels in HEK293 cells treated with DMSO control or GSK-A1 (100 nM, 24 h). (C) Live-cell imaging of PA localization in HEK293 cells expressing the PA biosensor GFP-Nir2-LNS2 (816-1181), treated with or without GSK-A1 (100 nM, 24 h) or with or without PLD inhibitor FIPI (1 μM). Scale bars: 10 µm. Quantification performed on five cells per condition. (D) Live-cell BRET assay analysis of PA produced at the PM on HEK293 cells treated with or without GSK-A1 (100 nM, 24 h) or with or without PLD inhibitor FIPI (1 μM) using the Nir2-LNS2 (816-1181) PA biosensor. (E) Schematic of IMPACT labeling for generating fluorescent reporters of PLD activity. (F) Normalized PLD activity by IMPACT on HEK293T cells treated with GSK-A1 (100 nM, 18 h) or DMSO with or without FIPI pretreatment (750 nM, 30 min). (G) Confocal microscopy imaging of the localization of PLD activity, assessed using real-time IMPACT (RT-IMPACT) labeling in HEK293 cells treated with GSK-A1 (100 nM, 24 h) and then either DMSO or FIPI (1 µM, 30 min). (H) Normalized PLD activity by IMPACT on HEK293T PLD1/2 double KO cells transiently overexpressing mScarlet-i empty vector or mScarlet-i-PLD1. Gray indicates untransfected population, and red indicates transfected population. (I) Normalized PLD activity by IMPACT on HEK293T cells treated with GSK-A1 (100 nM, 1 h), VT01454 (100 nM, 1 h), or DMSO with or without FIPI pretreatment (750 nM, 30 min). Different shapes on plot correspond to different biological replicates (n = 3). Statistical significance was determined using one-way ANOVA with Tukey post hoc test, with p values indicated on the plot.

Beyond DGKs, PA is produced by additional biosynthetic pathways in the cell, including the hydrolysis of PC by PLDs. Of the two PLD isoforms (PLD1/2) that produce PA via PC hydrolysis, PLD1 translocates to the PM to produce PA upon activation^45^ and PLD2 is constitutively localized at the PM^46^. We found that PA increases caused by GSK-A1 treatment at the PM were greatly reduced in cells pretreated with the selective pan-PLD1/2 inhibitor, FIPI^47^, as judged by confocal microscopy (Fig. 1C) or BRET analysis based on the Nir2-LNS2 (816-1181) biosensor (Figure 1D).

To confirm the involvement of PLDs, we used a small-molecule chemical biology tagging approach termed Imaging Phospholipase D Activity with Clickable Alcohols via Transphosphatidylation (IMPACT), which we previously developed to quantify PLD activity in single cells^48^. Briefly, IMPACT involves treatment of cells with an azide-containing primary alcohol, which is used as a substrate for PLD in a transphosphatidylation reaction that produces a phosphatidyl azidoalcohol lipid instead of PA. Next, cells are treated with a cyclooctyne-bearing fluorophore (e.g., BCN-BODIPY), which selectively tags the phosphatidyl azidoalcohols via a bioorthogonal reaction. The cells can then be analyzed by flow cytometry to quantify BODIPY-derived fluorescence as a readout of PLD activity (Figure 1E; see also Figure S1 for flow cytometry gating). IMPACT followed by flow cytometry revealed an increase in PLD activity of HEK293T cells treated with GSK-A1, and this increase was prevented by pretreatment with the PLD inhibitor FIPI (Figure 1F). Elevation of PLD1 levels using CRISPRa further augmented the GSK-A1- induced PLD activity (Figure S2B, C). We then used a real-time variant of IMPACT (RT-IMPACT) to visualize the subcellular localization of active PLDs^49^. Here, a different alcohol probe containing a *trans*-cyclooctene (TCO) is used along with fluorogenic tetrazine detection reagents in a bioorthogonal reaction termed the tetrazine ligation, which exhibits rapid kinetics and no-rinse imaging such that localizations of IMPACT-derived fluorescent lipids can be visualized prior to subsequent interorganelle transport. Using this RT-IMPACT protocol, we found that the localization of PLD activity induced by GSK-A1 was at the PM (Figure 1G). These experiments revealed that PLDs are involved in the GSK-A1-induced PA increase.

Further, in PLD1/2 double knockout HEK293T cells, overexpression of mScarlet-i-PLD1 but not mScarlet-i alone led to increased PA production following GSK-A1, verifying the specificity of the IMPACT readout to PLDs (Figure 1H). To further ensure that the PA increase was the result of PI4P loss rather than off-target effects of GSK-A1, we inhibited PM PI4P generation by interfering with PI transfer to the PM using the Class I PITP inhibitor VT01454, which prevents PI pools from the ER from being furnished to the PM, thus hindering PI4P synthesis^43^. We found that VT01454 treatment phenocopied GSK-A1, eliciting a comparable elevation of PLD activity in HEK293T cells (Figure 1I), with similar results observed in U2OS cells (Figure S3A). Together, these results demonstrate that disruption of PI4P synthesis at the PM triggers a response in which PLD activation drives a PA increase at the PM.

### PS counteracts PLD-mediated PA production under PI4P depletion

Because PLD-derived PA is not directly downstream of PI4P synthesis, we examined whether additional changes to lipid levels induced by PI4P depletion might be related to the increase of PA. Lipidomics analysis of HEK293 cells revealed a decrease in multiple PS species upon 24 h treatment with GSK-A1 (Figure 2A), suggesting that PI4P loss perturbs the pool of anionic lipids at the PM, consistent with our earlier work^42^. To determine whether PS abundance regulates PLD activation under PI4P depletion conditions, we performed IMPACT on cells overexpressing either wild-type phosphatidylserine synthase 1 (PSS1) or a gain-of-function mutant PSS1^P269S^ associated with Lenz-Majewski Syndrome that is insensitive to product inhibition and therefore causes substantial accumulation of PS in the cell^42^. We found that HEK293T cells overexpressing wild-type PSS1 exhibited a small decrease in the GSK-A1-induced PLD response, whereas those overexpressing PSS1^P269S^ showed almost no PLD response to GSK-A1 treatment (Figure 2B), with similar trends observed in U2OS cells (Figure S3B). By contrast, pharmacological inhibition of PS synthesis using a PSS1 inhibitor^50^ significantly amplified the PLD response to GSK-A1 (Figure 2D).

**Figure 2.**
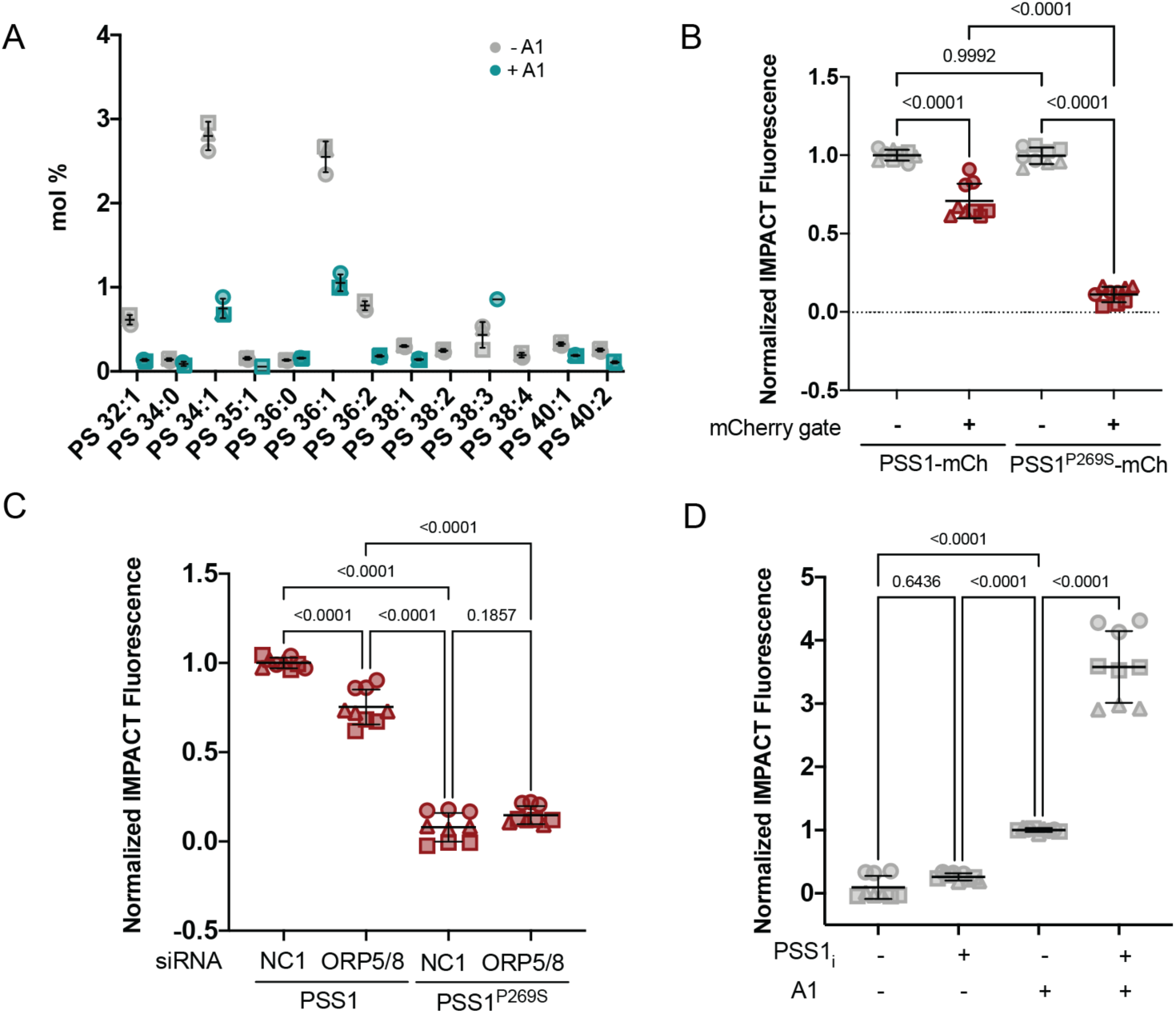
PS levels modulate the extent of PLD activation induced by PI4KIIIα inhibition. (A) Lipidomics analysis of PS levels in HEK293 cells treated with DMSO control or GSK-A1 (100 nM, 24 h). (B) Normalized PLD activity by IMPACT on HEK293T cells transiently overexpressing PSS1^WT^-mCherry or PSS1^P269S^-mCherry. Gray indicates untransfected population, and red indicates transfected population. (C) Normalized PLD activity by IMPACT on HEK293T cells knocked down by siRNA against ORP5 and ORP8 followed by transient overexpression of PSS1^WT^-mCherry or PSS1^P269S^-mCherry. (D) Normalized PLD activity by IMPACT on HEK293T cells pretreated with or without the PSS1 inhibitor DS55980254 (1 μM, 24 h) followed by treatment with or without GSK-A1 (100 nM, 4 h). Different shapes on plot correspond to different biological replicates (n = 3). Statistical significance was determined using one-way ANOVA with Tukey post hoc test, with p values indicated on the plot.

Because PS is delivered to the PM in exchange for PI4P via ORP5/8 transporters^11,12^, we next investigated whether such PI4P/PS counter-transport played a role in this phenomenon. Knockdown of ORP5/8 by siRNA caused only a minor change in PLD activity under control conditions and no effect at all under conditions of excess PS synthesis in the ER, i.e., overexpression of PSS1^P269S^ (Figure 2C). These results suggest that ORP5/8 transport does not play a pivotal role in the PI4P–PS–PA regulation and that other PS-related regulators are likely also involved. Nevertheless, these data reveal a reciprocal relationship between two negatively charged phospholipids, PA and PS, under conditions when levels of PI4P at the PM are reduced by PI4KIIIα inhibition. That is, the decreased PS synthesis induced by PM PI4P depletion, as a likely result of decreased PI4P-mediated PS transport, leads to elevation of PA at the PM, and modulation of PS levels under limited PI4P supply at the PM has an opposite effect on PLD-mediated PA synthesis.

### GSK-A1-induced PLD activation is mechanistically distinct from PKC-mediated PLD activation

Given the strong response of PLD activation to PI4KIIIα inhibition, we sought to establish mechanisms underlying this phenomenon. We first compared PI4P depletion-driven PLD activation via GSK-A1 with classical PKC-dependent PLC stimulation by measuring PLD activity in cells treated with GSK-A1 over an 18-h timecourse or via acute (30 min) treatment with the PKC activator phorbol 12-myristate 13-acetate (PMA). PMA, a chemical mimic of DAG, activates classical PKCs including protein kinase Cα (PKCα), which in turn acutely activate PLDs^51^. PMA triggered a rapid, maximal PLD response within minutes, whereas GSK-A1 elicited a slowly developing increase that plateaued after 4 h (Figure 3A and Figure S3C). We therefore hypothesized that PLD activation by PI4KIIIα inhibition might occur via a route that involves gene transcription and protein expression, which would match the slower timescale.

**Figure 3.**
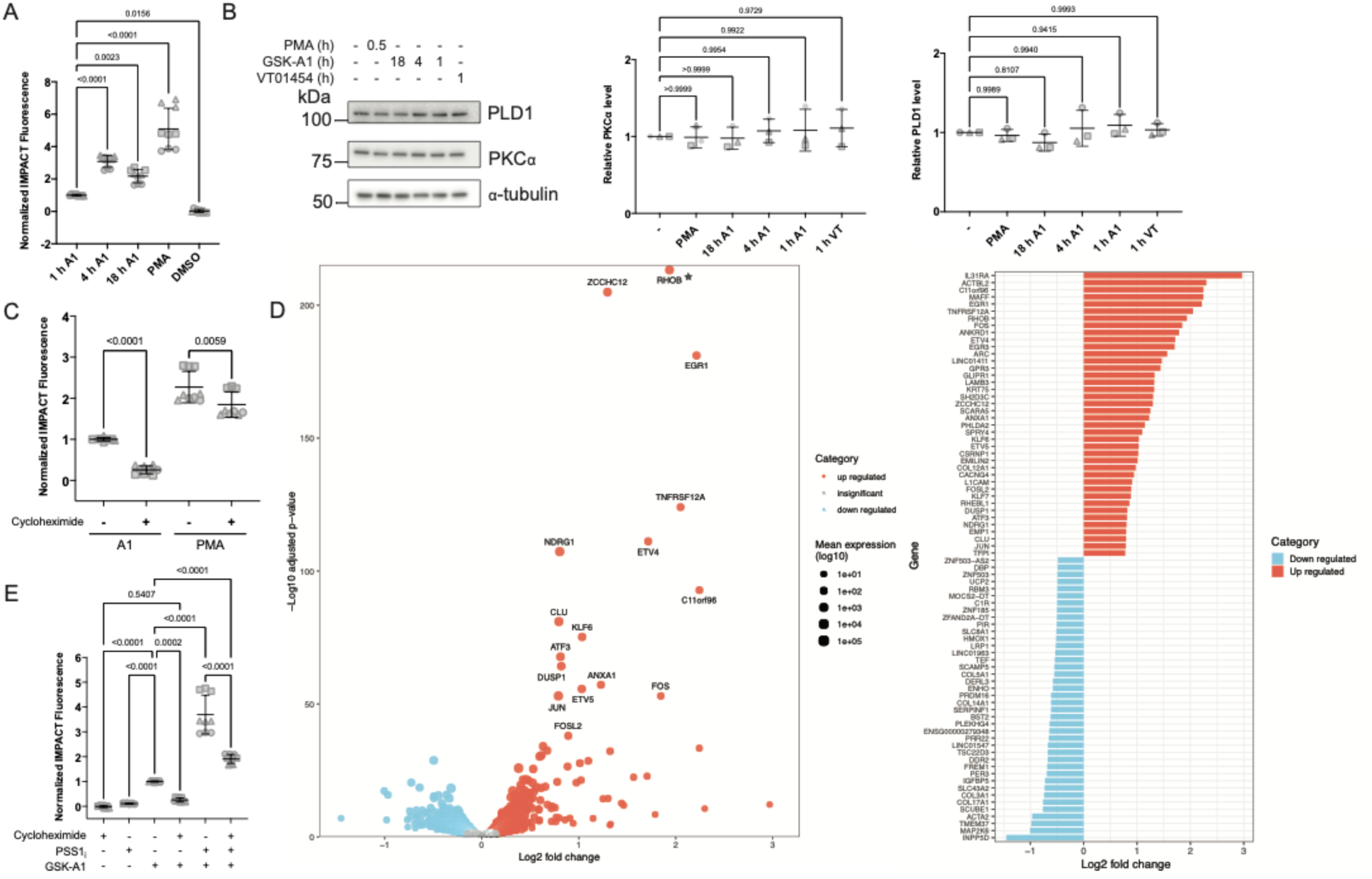
Depletion of PI4P at the plasma membrane leads to transcriptional and translational changes. (A) Normalized PLD activity by IMPACT on HEK293T cells treated with GSK-A1 (100 nM) for 0, 1, 4, or 18 h or with PMA (100 nM) for 30 min. (B) Western blot analysis of HEK293T cells treated with GSK-A1 (100 nM) for 0, 1, 4, or 18 h, PMA (100 nM) for 30 min, or VT01454 (100 nM) for 1 h. Quantification performed by normalization of PLD1 or PKCα protein band intensity to total protein content (Ponceau S). (C) Normalized PLD activity by IMPACT on HEK293T cells pretreated with cycloheximide (5 μg/mL, 1 h) followed by treatment with GSK-A1 (100 nM, 4 h) or PMA (100 nM, 30 min). (D) mRNA sequencing results on HEK293 cells treated with DMSO control or GSK-A1 (100 nM, 24 h). Volcano plot highlighting differentially expressed genes in A1 vs. control. The plot summarizes the GSK-A1 inhibitor treatment effect where points represent genes (n = 14,038). The x-axis shows the estimated log_2_ fold-change (A1 − control) obtained with adaptive shrinkage, and the y-axis shows –log_10_ of the Benjamini–Hochberg adjusted p-value. Genes colored red passed the 10% FDR threshold and have positive log_2_ fold-changes; genes colored blue passed the 10% FDR threshold and have negative log_2_ fold-changes; grey points are non-significant. Positive log_2_ fold-change values indicate higher expression in A1 treated cells, negative values indicate lower expression. Point size is determined by log_10_ expression across all samples (A1 and control) for the gene. The adjusted p-value for RHOB rounds to zero, and its −log10 value is infinity, which cannot be plotted. Its location was manually and arbitrarily set to the y-axis limit for the purpose of visualization. The horizontal bar chart at right displays the 40 most strongly up- and down- regulated genes by GSK-A1 inhibitor treatment that met the 10% FDR threshold. The x-axis shows the estimated log_2_ fold-change (GSK-A1 − control) derived with adaptive shrinkage; red bars extending right denote higher expression in A1-treated cells, while blue bars extending left denote lower expression. Gene symbols appear on the y-axis, ordered by descending effect size. Full dataset is provided in **Table S1**. (E) Normalized PLD activity by IMPACT on HEK293T cells treated with or without the PSS1 inhibitor DS55980254 (1 μM, 24 h), followed by treatment with or without cycloheximide (5 μg/mL, 1 h), followed by treatment with or without GSK-A1 (100 nM, 4 h). Different shapes on plot correspond to different biological replicates (n = 3). Statistical significance was determined using one-way ANOVA with Tukey post hoc test, with p values indicated on the plot.

We found no increase in PLD1 protein levels upon GSK-A1 treatment; in fact, prolonged treatment caused some degradation of PLD1 (Figure 3B and Figure S3D). We then treated cells with cycloheximide prior to GSK-A1 treatment to block protein synthesis and found that the GSK- A1-mediated effect on PLD activity was strongly reliant on *de novo* protein synthesis, in contrast to PMA treatment, whose effects on PLD activity were minimally changed by cycloheximide (Figure 3C). These results indicate that PI4P depletion engages a translation-dependent (and possibly transcription-dependent) pathway to activate PLD. Therefore, we then performed mRNA sequencing (RNA-seq) analysis of cells treated with GSK-A1 for 24 h (Figure 3D). These studies revealed upregulated mRNA levels of several transcription factors, including ZCCHC12, EGR1, ETV4, ETV5, KLF6, FOS, and JUN, as well as strong upregulation of the small GTPase RhoB following GSK-A1 treatment. To confirm that the upregulated mRNA levels were related to an increase in *de novo* protein synthesis, we additionally performed ribosome profiling (Ribo-seq) on cells treated with GSK-A1 for 0, 1, and 6 h and indeed found that RhoB exhibited upregulated translation upon GSK-A1 treatment (Figure S4).

To dissect the interplay between PS-mediated suppression and transcriptional control of PLD activation, we combined cycloheximide treatment with pharmacological PSS1 inhibition. Cycloheximide attenuated but did not abolish the enhanced PLD activation seen upon PS depletion, and the overall PLD activity remained above levels induced by GSK-A1 alone (Figure 3E). Thus, these data collectively support a model wherein the response to PM PI4P depletion involves a transcriptional response resulting in elevated PLD-mediated PA synthesis.

### RhoB enhances PLD activation in response to PI4P depletion

As described above, transcriptomics and Ribo-seq analyses revealed the upregulation of RhoB upon PI4KIIIα inhibition. RhoB shares 85% amino acid sequence identity with the known PLD activator RhoA^52^, but RhoB has not been previously implicated in PLD1/2 regulation. Consistent with the RNA-seq and Ribo-seq data, immunoblotting showed a substantial increase in RhoB protein levels after GSK-A1 treatment in both HEK293T and U2OS cells, with a more pronounced increase in U2OS cells, corresponding to their higher PLD activity response (Figures 4A and 3A). To assess whether RhoB is required for PI4P-depletion-driven PLD activation, we generated RhoB CRISPR knockout (KO) cell lines. As expected, these cells failed to upregulate RhoB protein levels upon GSK-A1 treatment, and we found that they exhibited a substantial reduction in GSK-A1-induced PLD activity compared to WT controls (Figures 4B–C). Conversely, CRISPRa-mediated increase of RhoB expression enhanced the levels of PLD activity in response to GSK-A1 (Figure 4D). We noted that previous studies have reported that tagging RhoB with fluorescent proteins can affect its localization and function^53,54^. Therefore, to transiently overexpress RhoB via plasmid transfection, we employed a bicistronic mScarlet-i-P2A-RhoB construct resulting in co-expression of essentially untagged RhoB and free mScarlet-i, minimizing the perturbation to RhoB while allowing for identification of transfected cells by flow cytometry. By performing IMPACT labeling on such cells overexpressing RhoB, we found an enhancement in GSK-A1-induced PLD activity compared to GSK-A1 alone (Figure 4E).

**Figure 4.**
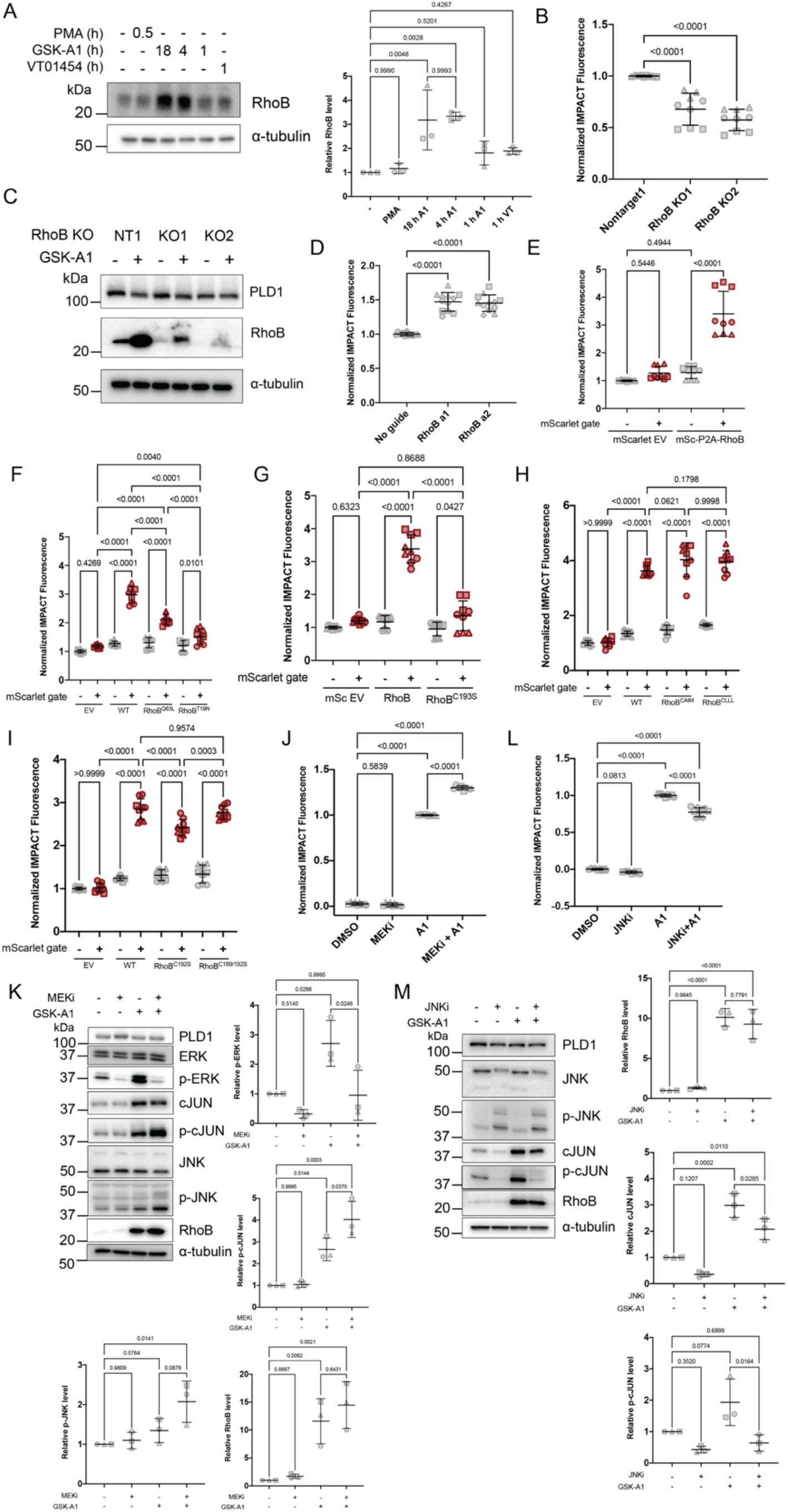
RhoB enhances PLD activity under conditions of plasma membrane PI4P depletion. (A) Western blot analysis of HEK293T cells treated with GSK-A1 (100 nM, 0–18 h), PMA (100 nM, 30 min), or VT01454 (100 nM, 1 h). Quantification performed by normalization of RhoB protein band intensity to total protein content (Ponceau S). (B) Normalized PLD activity by IMPACT on HeLa cells expressing Cas9 and the indicated sgRNA (RhoB or non-targeting) treated with or without GSK-A1 (100 nM, 18 h). (C) Western blot analysis of HeLa cells expressing Cas9 and the indicated sgRNA (RhoB or non-targeting) treated with or without GSK-A1 (100 nM, 18 h). (D) Normalized PLD activity by IMPACT on K562 CRISPRa cells expressing the indicated sgRNA (non-targeting or RhoB) treated with or without GSK-A1 (100 nM, 18 h). See Figure S6A for validation of RhoB CRISPRa. (E) Normalized PLD activity by IMPACT on HEK293T cells transiently overexpressing mScarlet-i empty vector or mScarlet-i-P2A-RhoB. Gray indicates untransfected population, and red indicates transfected population. (F-I) Normalized PLD activity by IMPACT on HEK293T cells transiently overexpressing mScarlet-i empty vector, mScarlet-i- P2A-RhoB, mScarlet-i-P2A-RhoB^Q63L^, mScarlet-i-P2A-RhoB^T19N^, mScarlet-i-P2A-RhoB^CAIM^, mScarlet-i- P2A-RhoB^CLLL^, mScarlet-i-P2A-RhoB^C193S^, mScarlet-i-P2A-RhoB^C192S^, or mScarlet-i-P2A-RhoB^C189/192S^ treated with or without GSK-A1 (100 nM, 18 h). Gray indicates untransfected population, and red indicates transfected population. (J) Normalized PLD activity by IMPACT on U2OS cells pretreated with or without the MEK inhibitor AZD6244 (2 μM, 24 h) followed by treatment with or without GSK-A1 (100 nM, 4 h). (K) Western blot analysis of U2OS cells pretreated with or without the MEK inhibitor AZD6244 (2 μM, 24 h) followed by treatment with or without GSK-A1 (100 nM, 4 h). Quantification performed by normalization of target protein band intensity to total protein content (Ponceau S). (L) Normalized PLD activity by IMPACT on U2OS cells pretreated with or without the JNK inhibitor JNK-IN-8 (1 μM, 24 h) followed by treatment with or without GSK-A1 (100 nM, 4 h). (K) Western blot analysis of U2OS cells pretreated with or without the JNK inhibitor JNK-IN-8 (1 μM, 24 h) followed by treatment with or without GSK-A1 (100 nM, 4 h). Quantification performed by normalization of target protein band intensity to total protein content (Ponceau S). Different shapes on plot correspond to different biological replicates (n = 3). Statistical significance was determined using one-way ANOVA with Tukey post hoc test, with p values indicated on the plot.

Because Rho GTPases function via GTP/GDP cycling, we assessed the importance of nucleotide binding for the effects of RhoB on PLD. IMPACT labeling of cells expressing either the GTP-locked RhoB^Q63L^ or the GDP-locked RhoB^T19N^ mutant forms^55^ resulted in no further increase to PLD activity by GSK-A1 compared to GSK-A1 alone, suggesting that dynamic GTPase cycling is important for the enhancing effect of RhoB on PLD activity (Figure 4F). RhoB undergoes prenylation at C193, which can influence its membrane localization and other downstream effects^56^. Mutation of C193 nearly completely abolished its ability to enhance PLD activity (Figure 4G), whereas mutations to the C-terminal CAAX motif to control the length of the prenylation tag (e.g., farnesylation using CAIM vs. geranylgeranylation using CLLL^57^) had equivalent stimulatory effects (Figure 4H). These results indicate that prenylation of RhoB was required for its PLD enhancement but that the form of prenylation did not impact this effect. Unlike RhoA, RhoB is also *S-*acylated (palmitoylated) adjacent to the prenylation site, and because palmitoylation can affect peripheral membrane protein localization and behavior within the bilayer^58^, we also investigated whether RhoB palmitoylation was important for its PLD enhancement activity. These studies revealed that ablation of the two C-terminal palmitoylation sites (C189 or C192) caused only a modest reduction in RhoB enhancement of PLD activity (Figure 4I), suggesting that palmitoylation fine-tunes but is not absolutely required for the enhancing effect of RhoB on PLD activity upon PI4KIIIα inhibition.

We next explored how PI4P depletion triggers RhoB upregulation. Among the transcription factors induced by GSK-A1 were several MAPK pathway regulators (Figure 3D and Figure S4) and c-Jun (Figure 3D). Based on previous studies reporting an association of RhoB with MAPK^59,60^, we used the MEK inhibitor AZD6244 to assess the involvement of MAPK pathways in the RhoB enhancement of PLD activity upon PI4P depletion. We found that AZD6244 treatment led to a small but significant increase in GSK-A1-induced PLD activity, accompanied by increased levels of RhoB, and phosphorylated forms of c-Jun and its kinase JNK (Figure 4J–K). PLD activity did not correlate with levels of phosphorylation of ERK1/2, suggesting that the p42/44 MAPK pathway is not involved in activation of PLD activity (Figure 4K). Interestingly, a prior study found that RhoB transcription is activated by c-Jun^60^. Therefore, to further examine potential roles of c-Jun and JNK involvement in RhoB expression and subsequent PLD activation upon PI4KIIIα inhibition, we treated cells with the JNK inhibitor JNK-IN-8 and found that it significantly blunted the GSK-A1-mediated activation of PLDs as well as levels of RhoB and phospho-c-Jun (Figure 4L–M). Overall, these data support a model wherein depletion of PI4P at the PM activates a c-Jun-dependent transcriptional program that contributes to RhoB upregulation, which amplifies PLD activity via its GTPase activity and membrane targeting to increase PLD-mediated PA synthesis.

### GSK-A1-stimulated PLD activity increases actin fiber formation

To gain additional information on how cells respond to PI4P depletion from the PM, including possible downstream mechanistic changes, we leveraged proximity labeling with a PM-anchored TurboID to assess proteomic changes in the local membrane environment upon PI4KIIIα inhibition (Figure 5A). To identify proteins recruited to the PM upon PI4P depletion, we performed such proximity labeling in HEK293T and HeLa cells treated with GSK-A1 or the PITP inhibitor VT01454, respectively (Figure 5B and Figure S5). Collectively, these studies revealed enrichment of multiple actin cytoskeleton-associated factors, including the Rho GTPase-binding protein Rhophilin-2 (RHPN2), which was previously implicated in RhoA and RhoB signaling and actin organization^61^. Supporting the specificity of this approach, we found that the protein that was most depleted from the PM upon PI4P inhibition was ORP8 (OSPBL8), consistent with its requirement of binding to PI4P in the PM for lipid transport functions^11,12^.

**Figure 5.**
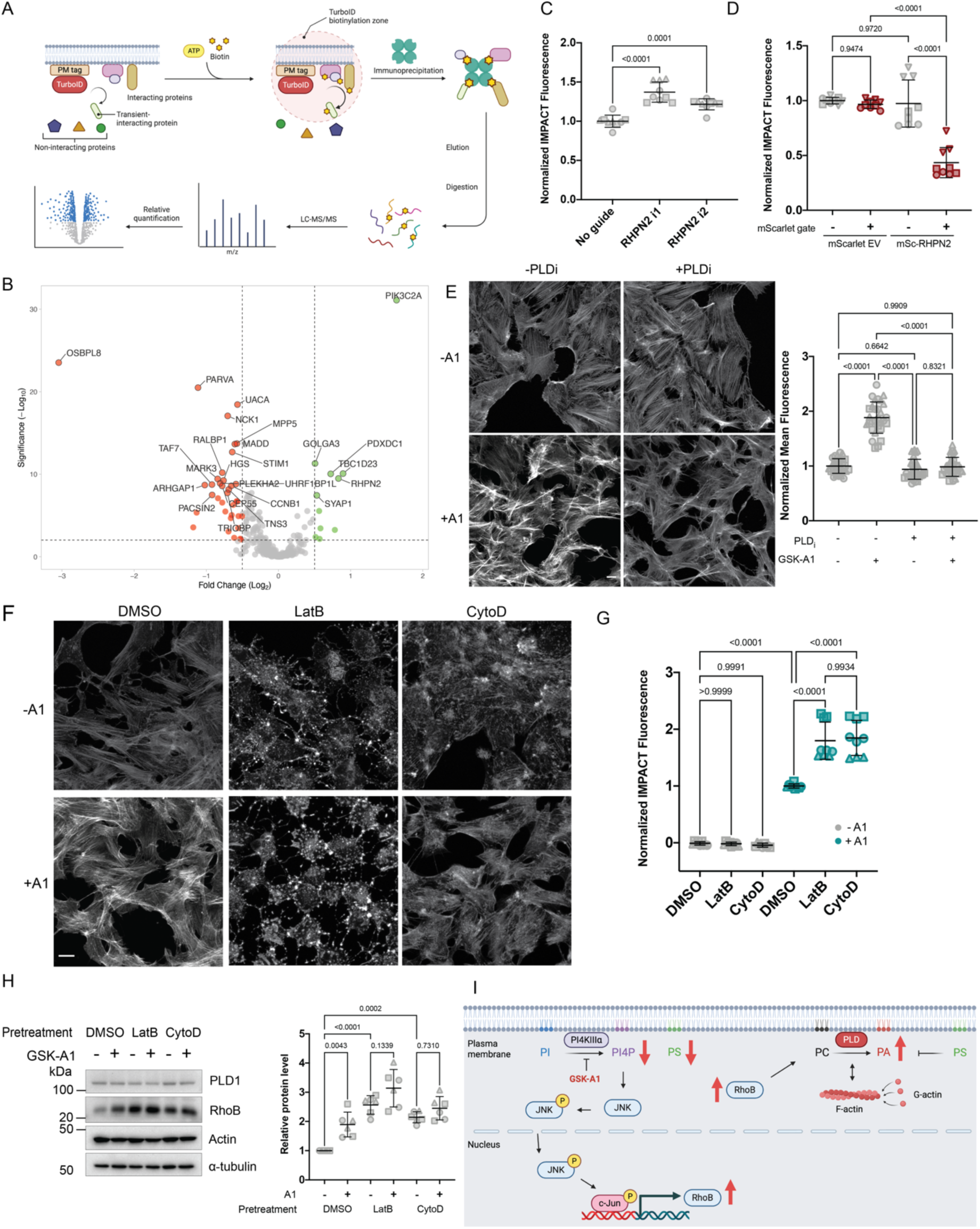
Acute PI4P depletion remodels actin cytoskeleton through PLD activity. (A) Schematic of plasma membrane-tagged TurboID proximity labeling proteomics. (B) TurboID proteomics results on HEK293T Lyn_10_-TurboID-V5 cells treated with or without VT01454 (100 nM, 1 h). Green indicates hits enriched and red indicated hits depleted from the PM after VT01454 treatment. Fold change in abundance is indicated on the x axis (in log_2_) and significance was determined using adjusted p values, shown on the y axis (in 1og_10_). Full dataset is provided in **Table S3**. (C) Normalized PLD activity by IMPACT on K562 CRISPRi cells with no sgRNA or RHPN2 sgRNAs treated with or without GSK-A1 (100 nM, 4 h). See Figure S6B for validation of RHPN2 CRISPRi cell lines using qPCR. (D) Normalized PLD activity by IMPACT on HEK293T cells transiently overexpressing mScarlet-i empty vector or mScarlet-i-RHPN2. Gray indicates untransfected population, and red indicates transfected population. (E) Phalloidin staining on U2OS cells pretreated with or without the PLD inhibitor FIPI (750 nM, 30 min) followed by treatment with or without GSK-A1 (100 nM, 4 h). Scale bar: 10 µm. Quantification performed by selecting 10 areas per condition over three frames per biological replicate. (F) Phalloidin staining on U2OS cells pretreated with or without Latrunculin B (2.5 μM, 1 h) or cytochalasin D (2.5 μM, 1 h) followed by treatment with or without GSK-A1 (100 nM, 4 h). Scale bar: 10 µm. (G) Normalized PLD activity by IMPACT on HEK293T cells pretreated with or without Latrunculin B (2.5 μM, 1 h) or cytochalasin D (2.5 μM, 1 h) followed by treatment with or without GSK-A1 (100 nM, 4 h). (H) Western blot analysis on on HEK293T cells pretreated with or without Latrunculin B (2.5 μM, 1 h) or cytochalasin D (2.5 μM, 1 h) followed by treatment with or without GSK-A1 (100 nM, 4 h). Quantification performed by normalizing RhoB protein band intensity over Ponceau S intensity. (I) Working model. Different shapes on plot correspond to different biological replicates (n = 3). Statistical significance was determined using one-way ANOVA with Tukey post hoc test, with p values indicated on the plot.

We next tested the role of RHPN2 in PLD regulation by using CRISPR interference. CRISPRi-mediated RHPN2 knockdown modestly but significantly increased GSK-A1-induced PLD activity, whereas RHPN2 overexpression had the opposite effect on PLD activity (Figure 5C– D), suggesting that RHPN2 alone can negatively regulate the PLD response to PI4KIIIα inhibition. Because RHPN2 can disassemble actin fibers when overexpressed^61^, we examined actin cytoskeleton organization after PI4P depletion. Using phalloidin staining, we found that GSK-A1 treatment induced a substantial increase in actin fiber levels, and importantly, this increase could be blocked by PLD inhibition (Figure 5E). Together with the enrichment of several actin-binding proteins at the PM upon PI4P depletion in our proximity proteomics datasets (Figures 5B and S5), these data suggest a causal relationship between PI4P depletion, PLD activation, and increased actin fiber formation.

Indeed, even under conditions that induce actin depolymerization, i.e., treatment with the actin-depolymerizing agents latrunculin B (LatB) or cytochalasin D (cytoD), PI4KIIIα inhibition partially rescued defects in actin fiber levels (Figure 5F). Further, treatment of under conditions of PI4KIIIα-inhibited cells with LatB and CytoD caused larger increases in PLD activity compared to PI4KIIIα inhibition alone, consistent with a role for LatB and CytoD in depleting G-actin, which can antagonize PLD activity^62^ (Figure 5G). Finally, the actin-depolymerizing agents also induced increases in RhoB protein levels, similar to effects of PI4KIIIα inhibition (Figure 5H). Overall, these data support a model (Figure 5I) wherein cells respond to lipidomic changes induced by PI4P depletion by activating a cascade culminating in RhoB expression, PLD-mediated PA synthesis, and F-actin assembly.

## DISCUSSION

In this study, we delineate a multicomponent integrated response that is activated by depletion of PI4P at the PM leading to stimulation of PLD enzymes for production of PA in this membrane. An important effect of the loss of PM PI4P is the concomitant decrease in total cellular PS levels. Under such conditions of low PM PI4P, levels of the anionic lipid PA are sensitive and reciprocally related to those of PS, the most abundant anionic lipid in the PM at steady state. These lipidomic changes are part of a series of molecular events that involve a complex set of transcriptional changes that include upregulation of the small GTPase RhoB, which enhances PLD activity and subsequently also promotes cortical actin fiber formation. Collectively, these findings reveal how cells integrate lipid metabolism and gene regulatory pathways to adjust membrane lipid levels in attempts to achieve homeostasis.

The negative charge of the PM inner leaflet, conferred by lipids such as PI4P, PS, and PA, is critical for recruitment and function of a number of peripheral proteins^7,63–65^. PI4P depletion would be expected to compromise this electrostatic landscape, potentially disrupting downstream signaling^7^. Rather than relying solely on PI(4,5)P_2_ buffering, cells appear to enlist PA synthesis as an alternative anionic lipid reservoir. PS and PA coexist as major anionic lipids at the PM, yet their relative abundances and distributions are dynamically interconverted via lipid-exchange proteins^66^. By manipulating PS synthesis and interorganelle transport, we reveal that PS and PA, two negatively charged lipids, may function in a rheostat to maintain PM inner leaflet charge and signaling competence under lipid stress conditions. In this model, PS occupies a gatekeeping function where when it is abundant, it suppresses PLD, preventing PA accumulation, but when it is scarce, PLD can be activated to produce PA.

Moreover, whereas GSK-A1 treatment alone did not stimulate progression through the classical PI cycle (e.g., as PLC activation would), it depletes a critical component, namely PI4P, that compromises the cell’s ability to generate second messengers when cell-surface receptors are activated. By invoking an alternate PA synthetic pathway, cells would provide the precursor for increased synthesis of PI as a compensatory response in an attempt to restore the decreasing PI4P levels in the PM.

The delayed kinetics of PLD-mediated PA production following PI4P inhibition hinted at a requirement for *de novo* protein synthesis. Whereas the transcriptional and translational responses to PI4KIIIα inhibition were indeed complex and multifaceted, important themes emerged that were bolstered by targeted mechanistic studies. One was the involvement of *de novo* gene expression via activation of the JNK/c-Jun pathway. A second was production of the small GTPase RhoB via this pathway and changes to a RhoB interactor RHPN2. A third was changes to organization of the actin cytoskeleton organization. Notably, overexpression of RhoB has been associated with increased actin fiber formation^67^, whereas overexpression of RHPN2 resulted in loss of actin fibers and cell contraction^61^. Co-overexpression of RhoB and RHPN2 caused increased actin stress fibers with a lack of cell contraction, suggesting that the interaction between these two proteins causes a distinct outcome from manipulating each protein in isolation. Our findings that increased actin fiber formation caused by PI4KIIIα inhibition was preventable by pretreatment with a PLD inhibitor, indicate that PA formation by PLD as a compensatory response to PI4P depletion directs increased actin fiber formation. These results are consistent with earlier work establishing that PLD-derived PA can stimulate F-actin fiber formation^68^ whereas free G-actin monomers negatively regulate PLD activity^62^. Our findings that perturbation with Latrunculin B or Cytochalasin D, both of which not only antagonize F-actin formation but also decrease levels of free G-actin monomers^69,70^, further increases PLD activity and RhoB upregulation upon PI4KIIIα inhibition, suggests a feedback loop wherein lipid perturbations can alter the actin cytoskeleton, in turn leading to further changes in the lipidome.

Actin cytoskeleton remodeling was one of the earliest phenotypes associated with perturbations to PI4P at the PM. In yeast, the PI4P pool produced by the PI4KIIIα ortholog Stt4 was shown to be crucial for vacuole morphology and actin cytoskeleton organization^9^. Further, the major PI4P-degrading enzyme is named suppressor of actin 1 (Sac1) because its mutation rescues a defective actin allele; further, its inactivation leads to an 8–10-fold increase in PI4P levels and can rescue defects in actin organization and growth caused by Stt4 mutants^9,71–75^. GSK-A1-treated mouse Schwann cells and the sciatic nerves of PI4KA KO mice both exhibit substantial actin remodeling, and conditional PI4KA KO mice also exhibit thinner myelin sheaths and defective nerve conduction^76^. Notably, defects in PI4KA function lead to congenital hypomyelinating leukodystrophy in humans, characterized by impaired myelin sheath formation in the central nervous system^35^. Myelination by highly specialized oligodendrocyte cells depends on extensive actin remodeling to extend and wrap PM processes around axons^77^. Our mechanistic studies here raise the possibility that for diseases associated with PI4KA dysfunction, insufficient PI4P production disrupts the PS–RhoB–PLD–PA circuit, leading to defective actin cytoskeletal organization at the leading edge of myelinating processes and resulting in hypomyelination.

In conclusion, in these studies we reveal how cells mount an integrated response to perturbations to PM PI4P pools. Lipidomic changes to PI4P and PS induce a reciprocal relationship between PS and PA in apparent efforts to bolster PM inner leaflet negative charge and boost the synthesis of PI, the main PI cycle intermediate, via stimulation of PA-synthesizing PLD enzymes. A key component of this response is *de novo* transcription of RhoB, whose activation of PLD-mediated PA synthesis not only causes lipidomic changes but also induces changes to F-actin organization. Our findings highlight the intricate crosstalk among lipid metabolic enzymes, small GTPases, and the actin cytoskeleton, providing a framework for understanding how PM lipids orchestrate both signaling and structural programs critical for cellular homeostasis.

## MATERIALS AND METHODS

### Antibodies

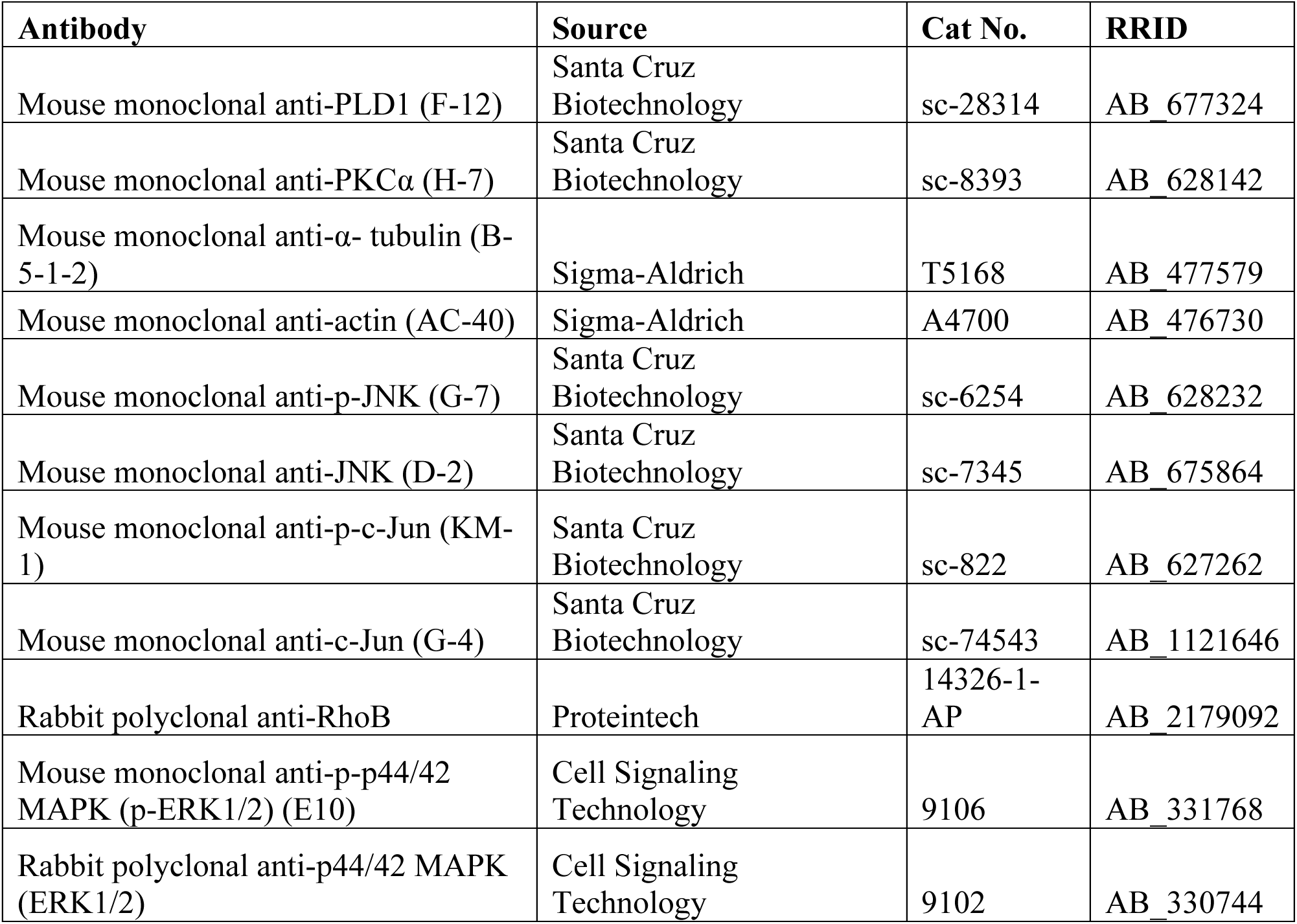

| Antibody | Source | Cat No. | RRID |
| --- | --- | --- | --- |
| Mouse monoclonal anti-PLD1 (F-12) | Santa Cruz Biotechnology | sc-28314 | AB_677324 |
| Mouse monoclonal anti-PKCα (H-7) | Santa Cruz Biotechnology | sc-8393 | AB_628142 |
| Mouse monoclonal anti-α- tubulin (B-5-1-2) | Sigma-Aldrich | T5168 | AB_477579 |
| Mouse monoclonal anti-actin (AC-40) | Sigma-Aldrich | A4700 | AB_476730 |
| Mouse monoclonal anti-p-JNK (G-7) | Santa Cruz Biotechnology | sc-6254 | AB_628232 |
| Mouse monoclonal anti-JNK (D-2) | Santa Cruz Biotechnology | sc-7345 | AB_675864 |
| Mouse monoclonal anti-p-c-Jun (KM-1) | Santa Cruz Biotechnology | sc-822 | AB_627262 |
| Mouse monoclonal anti-c-Jun (G-4) | Santa Cruz Biotechnology | sc-74543 | AB_1121646 |
| Rabbit polyclonal anti-RhoB | Proteintech | 14326-1-AP | AB_2179092 |
| Mouse monoclonal anti-p-p44/42 MAPK (p-ERK1/2) (E10) | Cell Signaling Technology | 9106 | AB_331768 |
| Rabbit polyclonal anti-p44/42 MAPK (ERK1/2) | Cell Signaling Technology | 9102 | AB_330744 |

### Reagents

Lipofectamine 2000, Invitrogen (11668019); Polybrene Infection/Transfection Reagent, Sigma-Aldrich (TR-1003); Trypsin Gold Mass Spectrometry Grade, Promega (V5280); D-biotin, Chem-Impex (00033, Cas #: 58-85-5); (+)-S-Trityl-L-cysteine, Alfa Aesar (L14384, CAS #: 2799- 07-7); Puromycin dihydrochloride, Sigma-Aldrich (P8833, CAS # 58-58-2); Doxycycline hyclate, Acros (446061000, CAS #: 24390-14-5); cOmplete Protease Inhibitor Cocktail, Roche (5056489001); GSK-A1, SYNkinase, (SYN-1219, CAS #: 1416334-69-4); Phorbol-12-Myristate-13-Acetate, Santa Cruz Biotechnology (sc-3576, CAS #: 16561-29-8); 5-gluoro-2-indolyl des-chlorohalopemide (FIPI), Cayman Chemical (13563, CAS #: 939055-18-2); JNK-IN-8, Ambeed Inc. (A206353, CAS #: 1410880-22-6); AZD6244, BioVision (2234-5, CAS #: 606143-52-6); Latrunculin B, Sigma-Aldrich (L5288, CAS #: 76343-94-7); Cytochalasin D, MedChemExpress (HY-N6682, CAS #: 22144-77-0); DS55980254, ProbeChem (PC-49548, CAS #: 2488609-41-0);32% Paraformaldehyde, Electron Microscopy Sciences (15714); Phalloidin-AF488, Invitrogen (A12379); Phalloidin-AF594, Invitrogen (A12381); Pierce™ High Capacity Streptavidin Agarose, Thermo Fisher (20359); Streptavidin-conjugated HRP, GeneTex (GTX85912); ProLong™ Diamond Antifade Mountant with DAPI, Thermo Fisher (P36971); Clarity™ Western ECL Substrate, Bio-Rad (1705061); BCA Protein Assay Kit, Thermo Fisher (23225).

The following compounds were prepared as described previously: 3-azido-1-propanol^78^; BCN-BODIPY^79^, VT01454^80^.

### Plasmids and cloning

mScarlet-i-tagged PLD1, PSS1-mCherry, and PSS1(P269S)-mCherry were generated previously^81,82^. The pSBtet-Lyn_10_-TurboID-V5-puro plasmid used for making a HeLa cell line stably expressing Lyn_10_-TurboID-V5 under doxycycline induction was generated previously^83^.

The PM-specific PI(4,5)P_2_ BRET biosensor, L_10_-mVenus-T2A-sLuc-PLCδ1_PH_ was generated previously^84^. To construct PM-specific PA BRET sensor, PLCδ1_PH_ domain was replaced with Nir2 LNS domain (residues 816-1181) from GFP-Nir2 (816-1181)^44^.

To construct the mScarlet-i-P2A-RhoB plasmid, the RhoB insert was amplified from pcDNA-GFP10-RhoB (Addgene 182237) and the mScarlet-i insert was amplified from the mScarlet-i-C1 vector (Addgene 85044). The mScarlet-i insert was subcloned into EcoRI/XhoI-digested pCAGGS-Lyn11-FRB*-dGFP-PLDs48-P2A-PLDs48-mCh-iFKBP (Masaaki Uematsu, Baskin Lab), replacing Lyn11-FRB*-dGFP-PLDs48. The RhoB insert was subcloned into KpnI/SalI-digested pCAGGS-mScarlet-P2A-PLDs48-mCh-iFKBP, replacing PLDs48-mCh- iFKBP. pCAGGS-mScarlet-i was made by inserting mScarlet-i into EcoRI/XmaI-digested M269. For amino acid mutagenesis, mutations were incorporated into the mScarlet-i-P2A-RhoB plasmid using the QuikChange XL Site-directed mutagenesis kit (Agilent).

To construct the mScarlet-i-RHPN2 plasmid, the RHPN2 insert was amplified from pDONR221-RHPN2 (DNASU 43557) and subcloned into the XhoI/KpnI-digested mScarlet-i-C1 vector.

For CRISPRa/i, sgRNAs against target genes were selected from the Weissman lab published dataset and cloned into the pCRISPRia-V2 vector using the BstXI and BlpI cut sites. For CRISPR KO, guides were designed with Benchling CRISPR tool and cloned into the BsmBI-digested pLentiCRISPR v2 vector.

### Primers

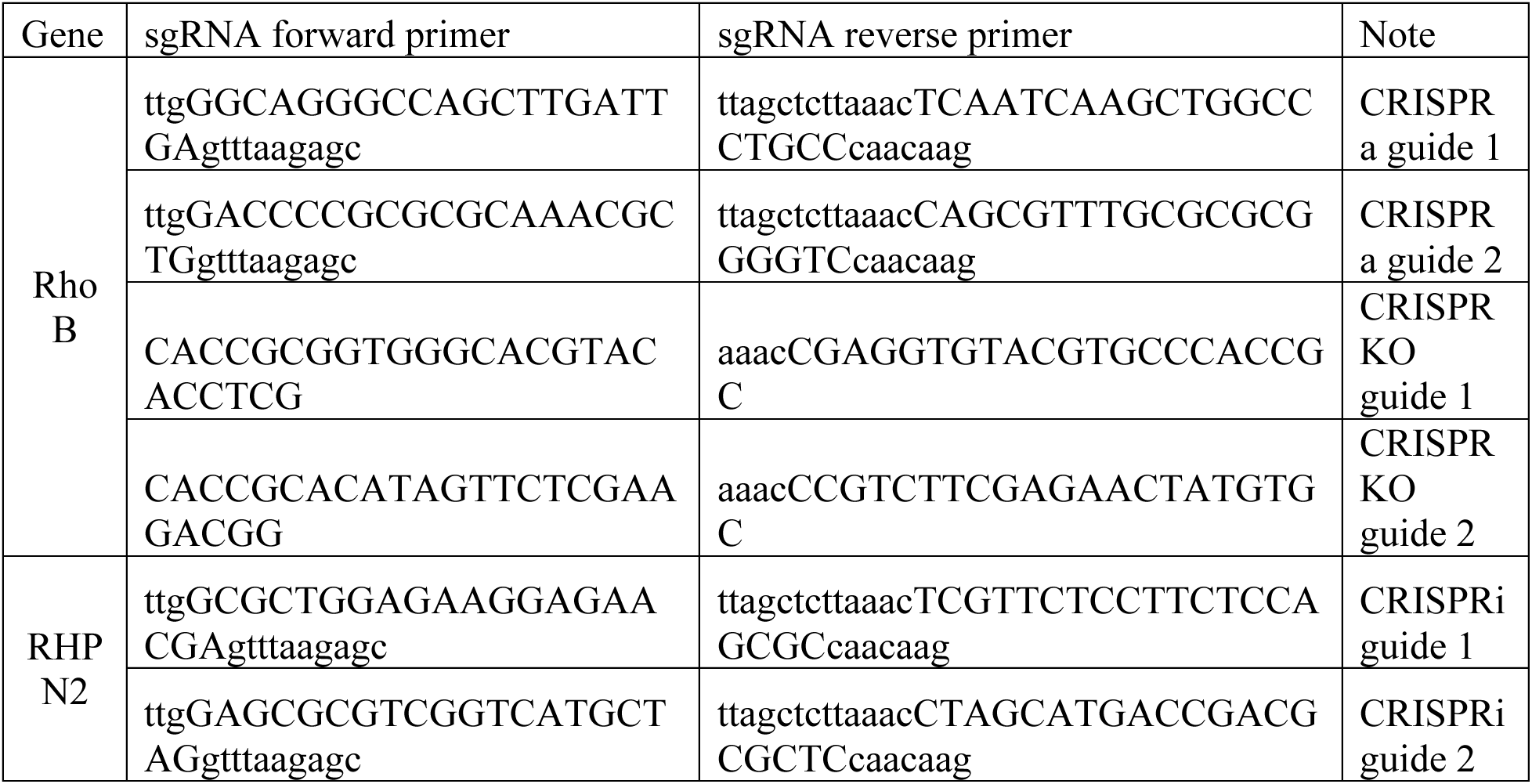
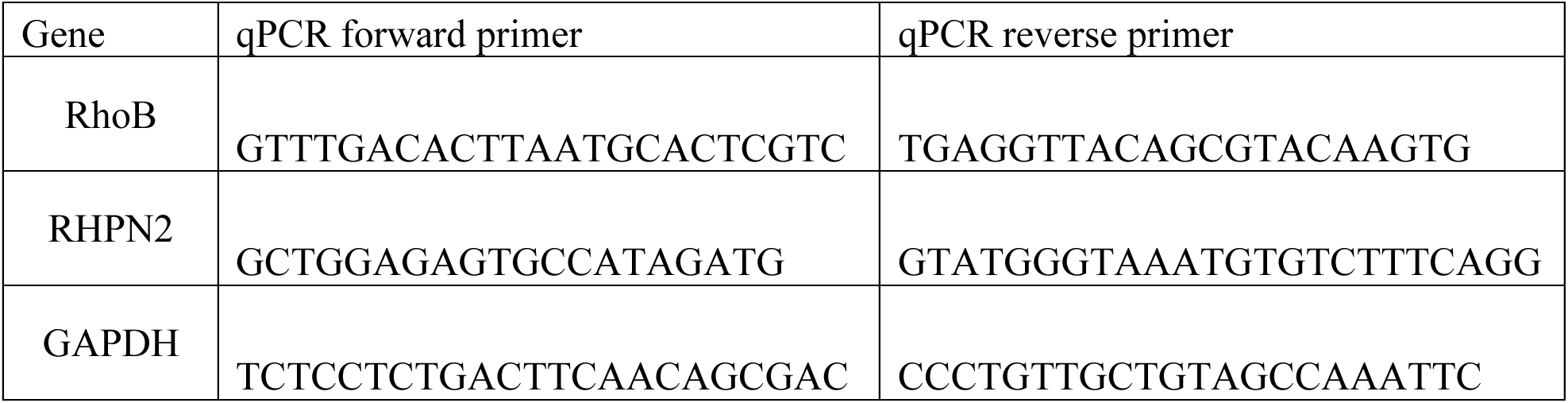

### Cell culture

All cells were maintained in a 5% CO_2_, water-saturated atmosphere at 37 °C. Flp-In T-Rex HeLa cells (Thermo Fisher) were maintained in DMEM (Corning) supplemented with 10% FBS (Corning) and 1% penicillin and streptomycin (P/S, Corning). HEK293T (ATCC) and HEK293TN (gift from Anthony Bretscher, Cornell University) cells were maintained in DMEM supplemented with 10% FBS, 1% P/S, and an additional 1 mM sodium pyruvate (Corning). K562-dCas9-KRAB (K562i) and K562-dCas9-VP64 cells (gifts from Martin Kampmann, UCSF) were maintained in RPMI 1650 medium (Corning) supplemented with 10% FBS, 1% P/S, and an additional 2 mM L-glutamine (Corning). U2OS cells were maintained in McCoy’s 5A medium (Corning) supplemented with 10% FBS and 1% P/S. Doxycycline-inducible lyn10-TurboID-V5 stable HeLa and HEK293T cell lines were generated in our previous studies^83^. HEK293T PLD1/2 double knockout cell line was previously generated^81^.

### Plasmid transfection

Plasmids were transfected into mammalian cells using 1 µL Lipofectamine 2000 : 2 μg DNA or 1 µL PEI : 3 μg DNA ratio following the manufacturer’s protocol. Cells were incubated with the transfection mix in Opti-MEM with 10% FBS for 6–8 h and then cultured in regular growth medium until harvest.

#### Lentivirus production

HEK293TN cells were seeded in 6-well plates with 2 mL media. Cells were transfected at 40-50% confluency. The transfection mixture was created by adding Lipofectamine 2000- containing Opti-MEM in 1:1 v/v ratio to DNA-containing Opti-MEM (2:1:3 Pax2:VSVg:lentiviral plasmid) and incubating for 20 min. Cell media was replaced with fresh Opti-MEM + 10% FBS and transfection mixture was added dropwise to HEK293TN cells. Opti-MEM was replaced with fresh growth medium after 6-8 h. Viral production was allowed to continue for 48 h with media harvested every 16 h. Virus-containing media was filtered through a 0.45 μm filter and store at 4 °C to be used within 1 week or aliquoted and stored in −80 °C for long-term storage.

#### Lentivirus transduction

For adherent cell lines, cells were seeded in 6-well plate with 2 mL media. One “kill control” well was included in which the cells would not be transduced with virus media. Cell media was replaced with 0.5 mL fresh media + 1.5 mL virus-containing media + 8 ug/mL polybrene at ∼60% cell confluency. Virus-containing media was replaced every 12 h for total of 3 times and cells were allowed to recover in fresh media for 12 h before antibiotic selection begins. For K562 suspension cells, 1 million cells were resuspended with 2 mL virus-containing media + 8 ug/mL polybrene and centrifuged at 1000 g for 2 h at 33 °C. After centrifugation, cells were recollected and centrifuged to removed virus-containing media, then resuspended with fresh RPMI media. Cells were allowed to recover overnight before antibiotic selection begins.

#### Validation

CRISPRa/i cell lines were validated by assessing mRNA levels by qPCR. CRISPR KO cell lines were validated by assessing protein levels by Western blot analysis.

### Confocal microscopy

For live-cell imaging, cells were seeded on 35 mm glass bottom dishes (14 mm diameter, #1.5 thickness, Matsunami Glass). For immunofluorescence, cells were seeded on 12 mm cover glass (#1.5 thickness, Fisherbrand) in 12-well plates (Corning). Glass bottom dish or cover glass was first incubated with 50 μg/mL poly-L-lysine in PBS for 2-5 h and rinsed three times with PBS before seeding of HEK293T cells. Cells were imaged live or fixed by immunofluorescence 24 h post transfection. Images were acquired via the Zeiss Zen Blue 2.3 software on Zeiss LSM 800 confocal laser scanning microscope equipped with Plan Apochromat objectives (40x 1.4 NA) and two GaAsP PMT detectors. Solid-state lasers (405, 488, and 561) were used to excite DAPI/tagBFP, EGFP/Alexa Fluor 488, and mCherry/mScarlet-i/Alexa Fluor 568, respectively. Acquired images were analyzed using FIJI.

#### Live-cell imaging

Cells were rinsed twice with PBS and imaged in Tyrode’s-HEPES buffer (T/H buffer: 135 mM NaCl, 5 mM KCl, 1.8 mM CaCl_2_, 1 mM MgCl_2_, 5 mg/mL glucose, 5 mg/mL bovine serum albumin, 20 mM HEPES, pH 7.4) at room temperature.

#### Immunofluorescence

Cells were rinsed twice with PBS, fixed in 4% paraformaldehyde in PBS for 10 min at room temperature, permeabilized with 0.5% Triton-X 100 in PBS for 5 min, and blocked in 1% BSA and 0.1% Tween-20 in PBS (blocking buffer) for 30 min. Cells were incubated with primary antibody in blocking buffer for 1 h at room temperature and rinsed three times with 0.1% Tween- 20 in PBS. Cells were then incubated with secondary antibody in blocking buffer for 1 h at room temperature in dark. Cells were rinsed three times with 0.1% Tween-20 in PBS before being mounted on slides in ProLong Diamond Antifade with DAPI and incubated overnight in dark before imaging. Slides were stored at 4 °C.

### Lipidomics analyses

Passage 6-9 of 600,000 HEK293 cells were plated onto 60 mm culture dishes and cultured overnight. The following day, GSK-A1 (100 nM) was added and cells were incubated for 24 h in DMEM medium with high glucose and 10 % serum. The harvested cell pellets were resuspended in PBS at a concentration of 6,000 cells/μL, then frozen and sent to Lipotype GmBH (Dresden, Germany) for mass spectrometry-based lipid analysis as described previously^85^. Briefly, lipids were extracted using a chloroform/methanol procedure^86^. Samples were spiked with internal lipid standard mixture. After extraction, the organic phase was transferred to an infusion plate and dried in a speed vacuum concentrator. The dry extract was resuspended in 7.5 mM ammonium formate in chloroform/methanol/propanol (1:2:4; v:v:v). All liquid handling steps were performed using Hamilton Robotics STARlet robotic platform with the Anti Droplet Control feature for organic solvents pipetting.

#### MS data acquisition

Samples were analyzed by direct infusion on a QExactive mass spectrometer (Thermo Scientific) equipped with a TriVersa NanoMate ion source (Advion Biosciences). Samples were analyzed in both positive and negative ion modes with a resolution of Rm/z=200=280000 for MS and Rm/z=200=17500 for MSMS experiments, in a single acquisition. MS/MS was triggered by an inclusion list encompassing corresponding MS mass ranges scanned in 1 Da increments (Surma et al. 2015).

#### Data analysis and post-processing

Data were analyzed with in-house developed lipid identification software based on LipidXplorer^87,88^. Data post-processing and normalization were performed using an in-house developed data management system. Only lipid identifications with a signal-to-noise ratio >5, and a signal intensity 5-fold higher than in corresponding blank samples were considered for further data analysis.

### Measurement of PM PI(4,5)P_2_ and PA using BRET-based sensors in live cells

HEK293 cells were seeded in white-bottom 96 well plates pre-coated with 0.01% poly-L-lysine solution (Sigma) and cultured overnight. Cells were then transfected with 0.1 μg of either the PI(4,5)P_2_ biosensor (L_10_-mVenus-T2A-sLuc- PLCδ1_PH_, for PI(4,5)P_2_ measurements) or the PM-targeted, Venus-tagged PA biosensor (Nir2-LNS2, residues 816-1181, for PA measurements) using Lipofectamine 2000 according to the manufacturer’s protocol. After 4-6 h of Lipofectamine incubation, the media was replaced with fresh media containing DMSO or GSK-A1 (100 nM), with or without FIPI (1 μM). After 24 h, the cells were quickly washed before being incubated for 30 minutes in 50 µL of modified Krebs-Ringer buffer at 37 °C in ambient air. After the pre-incubation period, the cell-permeable luciferase substrate, coelenterazine h (50 µL, final concentration 5 µM), was added and the signal from the mVenus fluorescence and sLuc luminescence were recorded using 485 and 530 nm emission filters over a 4 min baseline BRET measurement (15 s/cycle). The indicated inhibitors were maintained throughout all procedures. All measurements were performed in triplicate wells. BRET ratios (mVenus/Luciferase) were calculated for each well by dividing the 530-nm with the 485-nm intensity values. The processed BRET ratios obtained from drug-treated wells were then normalized to DMSO controls.

### mRNA sequencing

HEK293 cells were treated with DMSO control or GSK-A1 (100 nM) for 24 h. RNA was purified from frozen cells using RNeasy kits (Qiagen). Total RNA integrity was assessed on an Agilent 2100 Bioanalyzer RNA Nano chip and quantified by Qubit prior to library construction with the Illumina TruSeq Stranded RNA kit. Twelve libraries (three biological replicates, each with two technical replicates) were sequenced as 2 × 100 bp paired-end reads on an Illumina HiSeq 2500. lcdb-wf (https://github.com/lcdb/lcdb-wf) was used to perform all analysis steps up to and including differential expression analysis. Specifically, raw reads were adapter- and quality-trimmed with cutadapt v5.1^89^, then both pre- and post-trim QC metrics were collected with FastQC v0.12.1^90^ and aggregated using MultiQC v1.30^91^. Trimmed reads were aligned to GRCh38.p13 (GENCODE v46) using STAR v2.7.11b^92^, and gene-level counts were generated by featureCounts (Subread v2.1.1) using the GENCODE v46 GTF^93,94^. Differential expression analysis between A1 and control samples was conducted in R v4.2.3 using the DESeq2 v1.38.0 package^95^. Raw counts from technical replicates were summed for each biological replicate. A negative-binomial generalized linear model was fitted with the design formula ∼ treatment to contrast A1 versus control. Log2 fold-change estimates were shrunk using the adaptive shrinkage (ashr) method for interpretable effect-size estimation. To flag known transcription factors (TFs), we cross-referenced each gene against the curated set of 1,435 human DNA-binding TFs from Lovering et al.^96^ and added the binary transcription factor indicator to the results (Table S1). Enrichment of transcription factors among DEGs was assessed by Fisher’s exact test on a 2×2 contingency table comparing TF versus non-TF status and significant versus non-significant differential expression.

### Data availability

Raw FASTQ files and DESeq2 library–size normalized counts (counts.tsv) have been deposited in the Gene Expression Omnibus under accession GSE303723.

### Polysome profiling

Four plates (10-cm) of HEK293T cells/condition was grown to 80% confluency and treated with GSK-A1 (100 nM) for 0, 1, or 6 h. Cells were washed with cold PBS and lysed in the polysome lysis buffer (10 mM HEPES, pH 7.4, 100 mM KCl, 5 mM MgCl_2_, 100 μg/mL cycloheximide with 1% Triton X-100). The nuclei were pelleted by spinning at 21,130 g for 10 min at 4 °C. 500 µL of lysates were loaded onto a 15-45% (wt/vol) sucrose density gradients freshly prepared in a SW41 ultracentrifuge tube (Beckman) using a Gradient Master (BioComp Instruments). Samples were centrifuged at 180,000 g for 2.5 h at 4 °C in a Beckman SW41 rotor. Polysome profiles were recorded at A254 using the Brandel Gradient Fractionation System and an ISCO UA-6 UV/Vis detector.

#### Ribo-seq library construction

The cDNA library construction follows the Ezra-seq method described previously with minor modifications^97^. In brief, an aliquot of ribosome fractions representing monosome and polysome were collected followed by digestion with *E. coli* RNase I (Ambion, 750 U per 100 A260 units) by incubation at 4 °C for 1 h. RNA was extracted using Trizol LS reagent (Invitrogen) followed by ethanol precipitation. The ribosome-protected mRNA fragments (RPFs) were separated on a 15% polyacrylamide TBE-urea gel (Invitrogen) and visualized using SYBR Gold (Invitrogen). Selected regions in the gel corresponding to 25-35 nt were excised and dissolved by soaking in 400 µL RNA elution buffer (300 mM NaOAc pH 5.2, 1 mM EDTA, 0.1 U/µL SUPERase·In) at 4 °C for overnight. The gel debris was removed using a Spin-X column (Corning), followed by ethanol precipitation. 10∼200 ng RNAs were mixed with 10 U T4 PNK (NEB), 1 µL homemade Ezra enzyme, 5 U Poly(A) Polymerase (NEB), and 20 U SUPERase·In in Ezra buffer and incubated at 37 °C for 30 min followed by 70 °C for 10 min. Ligation was performed for 60 min at 25 °C by adding a 10 µL reaction mixture (0.5 μM biotinylated 5’ end adaptor, 1 × T4 Rnl2 reaction buffer, 20 U SUPERase·In, 15% PEG8000 and 10 U T4 RNA ligase 2 truncated KQ (NEB)). The ligated RNA sample was mixed with 10 µL of pre-washed streptavidin beads (NEB) and incubated at room temperature for 10 min. After washing once with 2 × SSC, beads were re- suspended in 12 µL nuclease-free water and mixed with 8 µL cDNA synthesis mixture (5 μM reverse transcription primer, 5 × first strand buffer, 0.1 M DTT, 10 mM dNTP, SuperScript III) followed by incubation at 50 °C for 30 min. After washing once with 2 × SSC, the cDNA was amplified by PCR using barcoded sequencing primers. PCR was performed by mixing 1 × HF buffer, 0.5 mM dNTP, 0.25 μM PCR primers and 0.025 U Phusion polymerase. PCR was carried out under the following conditions: 98 °C, 30 s; (98 °C, 5 s; 68 °C, 15 s; 72 °C, 20 s) for 14 cycles; 72 °C, 2 min. PCR products were separated on a 8% polyacrylamide TBE gel (Invitrogen). DNA products with the expected size around 180 bp were excised and recovered from DNA elution buffer (300 mM NaCl, 1 mM EDTA). After quantification by Agilent BioAnalyzer DNA 1000 assay, equal amounts of barcoded samples were pooled and sequenced using NextSeq 500 (Illumina).

#### Ribo-seq analysis

To align sequencing reads, the 5’ and 3’ adapters of the reads were trimmed by Cutadapt (version 2.8). The trimmed reads with length shorter than 15 nucleotides were excluded from the analysis. To keep accurate reading frame of Ribo-seq, low-quality bases at both ends of the reads were not subject to clip. The trimmed reads were first aligned to rRNAs using Bowtie (version 1.2.3)^98^. The rRNA sequences were downloaded from the nucleotide database of NCBI and RNAcentral. The reads unaligned to rRNAs were then mapped to the custom human transcriptome using STAR (version 2.7.10a)^92^. To avoid ambiguity, reads mapped to multiple positions or with >2 mismatches were disregarded for further analysis. The custom transcriptome was generated based on the reference genome and annotations obtained from Ensembl using human release 109 (GRCh38.p14). Protein coding genes were extracted, and a single transcript was selected for each gene on the following procedure. For each gene, the transcript with the longest coding sequence (CDS) was initially selected. If the selected transcripts have equal CDS length, the longest transcript was included in the custom transcriptome. Mapping to ribosomal P sites was performed by shifting the read position from the 5’ ends to the position by 12 nt.

#### Data and code availability

The sequencing data reported in this manuscript have been deposited in NCBI’s Gene Expression Omnibus under accession number GSE308521. Custom Python scripts used to analyze the sequencing data are available at GitHub and Zotero at https://github.com/usa0ri/Huang2025 and https://doi.org/10.5281/zenodo.17144598.

### Proximity biotinylation with Lyn_10_-TurboID

pSBtet-Lyn10-TurboID-V5 expression were induced in HeLa or HEK293T cells with 2.5 μg/mL doxycycline for 48 h, treated with 0, 1, 6 h GSK-A1 or 1 h VT01454, respectively, and then incubated with 500 μM biotin for 10 min at 37 °C under 5% CO_2_. Cells were rinsed five times with PBS. Cell pellet was collected by trypsinizing cells and rinsing three times with PBS at 500 g, 4 °C for 3 min. Cell pellet was resuspended with RIPA buffer (150 mM NaCl, 25 mM Tris pH 8.0, 1 mM EDTA, 0.5% sodium deoxycholate, 0.1% SDS, and 1% Triton X-100) supplemented with cOmplete protease inhibitor cocktail and sonicated using a tip sonicator at 20 % amplitude, 1 s on 1 s off, for 4 s. The cell mixture was centrifuged at 13,000 g for 5 min at 4 °C to clarify lysates. Protein concentration was quantified using the BCA assay (Thermo Fisher), and a small fraction of the clarified cell lysates was saved and normalized as input. The remaining cell lysate was subjected to pulldown using streptavidin-agarose with rotation at 4 °C overnight (12-16 h). The resin was then centrifuged for 5 min at 1000 g, washed two times with RIPA buffer, one time with 1 M KCl, once with 0.1 M Na_2_CO_3_, once with 2 M urea in 10 mM Tris pH 8.0, and twice with RIPA buffer to reduce non-specific binding. Samples were then denatured and analyzed by SDS-PAGE and Western blot with detection by chemiluminescence using Clarity Western ECL substrate.

#### TurboID proteomic sample preparation

Cells were grown in five to six 15-cm dishes per condition. Biotinylation and streptavidin IP were carried out as described above. Following the last RIPA rinse, the GFP-Trap magnetic resin was incubated with elution buffer (100 mM Tris pH 8.0, 1% SDS) at 95 °C for 5 min and the resin was spun at a tabletop centrifuge for 5 s and the supernatant was collected. This step was repeated two more times and supernatants were collected and combined for TMT labeling sample preparation. IP eluates were reduced by using 200 mM TCEP for 1 h at 55 °C. After that, samples were alkylated for 30 minutes at room temperature and in the dark using 375 mM iodoacetamide. Trypsin Gold, mass spectrometry grade (catalog no. V5280; Promega), was used to digest the samples at a ratio of 1:100 enzyme to substrate. The samples were then incubated to overnight at 37 °C. The Pierce Quantitative Colorimetric Peptide Assay (catalog no. 23275; Thermo Scientific) was utilized to quantify the concentrations of peptides. For TMT tests, samples were resuspended and normalized using 1 M triethylammonium bicarbonate (catalog no. 90114; Thermo Scientific). Samples were labeled using TMT 16plex label reagent sets (catalog no. A44520; Thermo Scientific) at a (w/w) label-to-peptide ratio of 20:1 for 1 h at room temperature. Labeling reactions were quenched by the addition of 5% hydroxylamine for 15min and pooled and dried using a SpeedVac. Labeled peptides were enriched and fractionated using Pierce High pH Reversed-Phase Peptide Fractionation Kit according to the manufacturer’s protocol (catalog no. 84868; Thermo Scientific). Liquid chromatography–tandem mass spectrometry fractions were analyzed using an EASY-nLC 1200 System (catalog no. LC140; Thermo Scientific) equipped with an in-house 3 μm C18 resin-(Michrom BioResources) packed capillary column (125 μm × 25 cm) coupled to an Orbitrap Fusion Lumos Tribrid Mass Spectrometer (catalog no. IQLAAEGAAPFADBMBHQ; Thermo Scientific). The mobile phase and elution gradient used for peptide separation were as follows: 0.1% formic acid in water as buffer A and 0.1% formic acid in 80% acetonitrile as buffer B; 0–5 min, 5%-8% B; 5–65 min, 8–45% B; 65–66 min, 45%-95% B; 66–80 min, 95% B; with a flow rate set to 300 nl min−1. MS1 precursors were detected at m/z = 375–1500 and resolution = 120,000. A CID-MS2-HCD-MS3 method was used for MSn data acquisition. Precursor ions with charge of 2+ to 7+ were selected for MS2 analysis at resolution = 50,000, isolation width = 0.7 m/z, maximum injection time = 50 ms and CID collision energy at 35%. 6 SPS precursors were selected for MS3 analysis and ions were fragmented using HCD collision energy at 65%. Spectra were recorded using Thermo Xcalibur Software v.4.4 (catalog no. OPTON-30965; Thermo Scientific) and Tune application v.3.4 (Thermo Scientific). Raw data and the V5 sequence (ggcaagcccatccccaaccccctgctgggcctggacagcacc) were searched using Proteome Discoverer Software 2.5 (Thermo Scientific) against an UniProtKB human database.

#### Downstream proteomic analysis

The computational tool Magma was used to analyze mass spectrometry proteomics data. Magma quantifies the differences in protein abundance between different experimental conditions by calculating fold-change (FC) and p-values using a customized linear mixed-effects model inspired by the MSStats TMT package^99,100^. Custom normalization was performed using the V5 sequence, assumed to remain constant between conditions. By comparing each bait protein against untransfected HeLa or HEK293T cells, the bait protein’s interactors were identified using criteria of fold change (FC) > 2, adjusted p-value < 0.05, and peptide-spectrum matches (PSM) > 5. Analysis was then narrowed to the combined set of interactors for both DMSO control and GSK-A1/VT01454-treated samples. To elucidate the specific effects of the GSK-A1/VT01454 treatment, we generated a volcano plot using the FC and adjusted p-values derived from the comparison between the GSK-A1/VT01454-treated samples and the DMSO control samples. Known contaminants in AP-MS experiments, including keratin (KRT), myosins (MYO), small ribosomal subunit proteins (RPS), heat shock-related 70 kDa proteins (HSPA), and large ribosomal subunit proteins (RPL), were excluded from the analysis.

### Western blot

Cells were lysed with RIPA buffer (150 mM NaCl, 25 mM Tris pH 8.0, 1 mM EDTA, 1% Triton X-100, 0.5% sodium deoxycholate, and 0.1% SDS) supplemented with cOmplete protease inhibitor cocktail on ice. Cell lysate was sonicated with a tip sonicator at 20% intensity, 1 s on 1 s off, for 4 s and centrifuged at 13,000 g for 5 min at 4 °C to clear lysate. Protein concentrations of the clarified cell lysates were quantified using the BCA assay. Cell lysates were normalized to 1– 5 mg/mL, and denatured with 6X Laemmli buffer (13.3% SDS, 0.067% bromophenol blue, 52.2% glycerol, 67 mM Tris pH 6.8 and 11.1% β-mercaptoethanol) at 95 °C for 5 min. Samples were analyzed by SDS-PAGE and Western blot with detection by chemiluminescence using Clarity Western ECL substrate (Bio-Rad).

### Agonists and inhibitors treatment summary

The following compounds were added to cells in conditions as summarized:

PLD activators: PMA, 100 nM, 30 min; GSK-A1, 100 nM, 1–24 h; VT01454, 100 nM, 1 h.

PLD inhibitor: FIPI, 750 nM, 30 min prior to PLD stimulation.

PSS1 inhibitor: DS, 1 μM, 24 h.

MEK inhibitor: AZD6244, 2 μM, 24 h.

JNK inhibitor: JNK-IN-8, 1 μM, 24 h.

Actin polymerization inhibitors: Latrunculin B, 2.5 μM, 1 h prior to PLD stimulation; Cytochalasin D, 2 μM, 1 h prior to PLD stimulation.

Protein expression inhibitor: cycloheximide, 50 μg/mL, 1 h prior to PLD stimulation.

### IMPACT labeling and flow cytometry

For adherent cell lines, cells were seeded in 24-well plate and allowed to grow for 2 days before transfection or drug treatment. After transfection or any drug pretreatment, negative control cells were treated with 750 nM FIPI for 30 min for subtraction of IMPACT background. Cells were subsequently treated with PLD stimulus for indicated time. Cells were then treated with 1 mM azidopropanol for 30 min at 37 °C to generate azido phosphatidylalcohols. Media was then aspirated, and cells were rinsed with three times with PBS. Cells were then incubated with prewarmed 1 μM BCN-BODIPY in T/H buffer (135 mM NaCl, 5 mM KCl, 1.8 mM CaCl_2_, 1 mM MgCl_2_, 5 mg/mL glucose, 5 mg/mL bovine serum albumin, 20 mM HEPES, pH 7.4) at 37 °C for 30 min, rinsed 2x with PBS, and incubated with prewarmed T/H buffer at 37 °C for 15 min to rinse out unreacted BCN-BODIPY. After two more PBS rinses, cells were trypsinized and transferred to a 96-well V-bottom plate and centrifuged at 1000 g, 4 °C to remove supernatant. Cells were resuspended in 175 µL PBS/well and centrifuged at 1000 g, 4 °C for rinsing. After two more rinses, cells were resuspended in 4% paraformaldehyde in PBS and incubated at room temperature in the dark for 10 min. After two more rinses with FACS buffer (PBS + 0.1% FBS), cells were resuspended with 120–150 µL of FACS buffer and subjected to flow cytometry analysis. A 488 nm laser was used to excite BODIPY-derived IMPACT fluorescence as a measure of PLD activity, and a 561 nm laser was used to excite the mScarlet-i-or mCherry-containing constructs. For experiments involving two colors, compensation was performed using single color control cells and unstained cells, and IMPACT fluorescence from cells with similar mScarlet-i/mCherry expression levels were extracted for data analysis. A BD Accuri instrument was used for 1-color flow cytometry and a Thermo Fisher Attune instrument was used for 2-color flow cytometry analysis. Cells were gated to exclude cell debris followed by singlet isolation, then FL2-A or FL1- A values were recorded. For each experimental condition, the average of median fluorescence intensities of the negative control FIPI-treated samples were subtracted from the respective no FIPI samples, and the differences of median fluorescence were normalized and plotted.

### RT-IMPACT labeling and confocal microscopy

HEK293 cells were labeled as previously described^49^. Briefly, cells (500,000) were seeded onto 29 mm glass-bottom dishes (CellVis, Cat# D29-20-1.5-N) pre-coated with 0.01% poly-L-lysine solution one day prior to imaging. All media used for labeling was pre-warmed to 37 °C. Cells were pretreated with the PLD inhibitor FIPI or DMSO vehicle control in culture media for 30 min at 37 °C. Following pretreatment, the media was completely aspirated, especially media within the glass well at the center of the dish. A 200 µL solution of 3 mM trans-5-oxocene (oxoTCO) was prepared in media, with or without FIPI, and was then added to cover only the central glass area of the dish and incubated for 10 min at 37 °C. After incubation, the oxoTCO solution was completely aspirated, followed by a brief rinse with 1 mL of fresh media and a final long rinse (5 min) with an additional 1 mL of fresh media at 37 °C. After rinsing, 1 mL of pre-warmed T/H buffer was added, and the dish was quickly placed on the microscopy stage and cells were put into focus. The buffer was then carefully removed while the dish was on the stage, and 100 µL of pre-warmed T/H buffer was added to the central glass well. Time-lapse imaging was initiated, followed by the addition of 100 µL of a 2 µM (2X) tetrazine-BODIPY solution in pre-warmed T/H buffer, resulting in a final concentration of tetrazine-BODIPY of 1 µM. Time-lapse imaging was commenced at 37 °C, with frames acquired every 3-4 s, followed by addition of Tz-BODIPY (1 µM). Images were acquired for the subsequent 1–2 min. Shown are images of the earliest timepoints with detectable fluorescent signal above background (∼10 s).

### Statistical analysis

For all experiments involving quantification, statistical significance was calculated in GraphPad Prism using tests as indicated in the figure legend. P values for each comparison are reported, and the number of biological replicates or cells analyzed is stated in the legend. All the raw data were plotted into graphs using GraphPad Prism. For all scatter plots, the black line indicates the mean.

## ACKNOWLEDGMENTS

We acknowledge support from the NIH (J.M.B.: R01GM151682, S.Q.: DP1GM142101). We thank members of the Baskin lab for helpful discussions. This research was supported in part by the Intramural Research Program of the National Institutes of Health (NIH). The contributions of the NIH author(s) are considered Works of the United States Government. The findings and conclusions presented in this paper are those of the author(s) and do not necessarily reflect the views of the NIH or the U.S. Department of Health and Human Services. Confocal imaging was performed in the Microscopy Core of NICHD with the kind assistance of Drs. Vincent Schram and Ling Yi.

## AUTHOR CONTRIBUTIONS

Conceptualization: S.H., Y.K., T.B., J.M.B.; Funding Acquisition: T.B., J.M.B.; Investigation: S.H., Y.K., X.C., T.W.B., M.S., M.T.M., R.D., J.K., S.U., S.G.; Project Administration: T.B., J.M.B.; Supervision: S.Q., H.Y., T.B., J.M.B.; Writing – original draft: S.H., J.M.B.; Writing – review & editing: S.H., Y.K., T.B., J.M.B.

