## Supplementary Information for "Phosphatidylserine and RhoB connect phosphatidylinositol 4-phosphate and phosphatidic acid metabolism at the plasma membrane"

**Contents:**

**Figures S1–S5**
**Legends for Tables S1–S3**

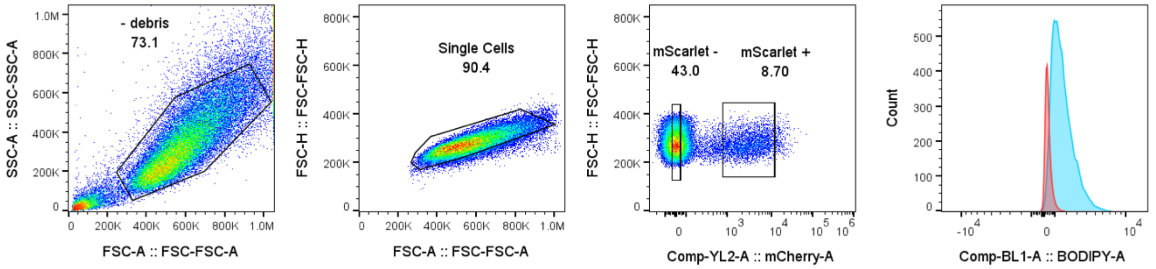


| Subset Name | Count | Median : Comp-BL1-A | Median : Comp-YL2-A |
|---|---|---|---|
| mScarlet + | 3160 | 337 | 2841 |
| mScarlet - | 15612 | 1466 | 1.89 |

**Figure S1. Flow cytometry sample gating strategy.** Cells were first plotted on FSC-A vs SSC-A to exclude debris. Subpopulation was then plotted on FSC-A vs FSC-H to exclude doublets. Single cells were plotted on YL-2A channel to be divided into mSc/mCh positive vs negative populations for two-color flow cytometry. Cells were plotted on BL1-A histogram for BODIPY median fluorescence statistics.

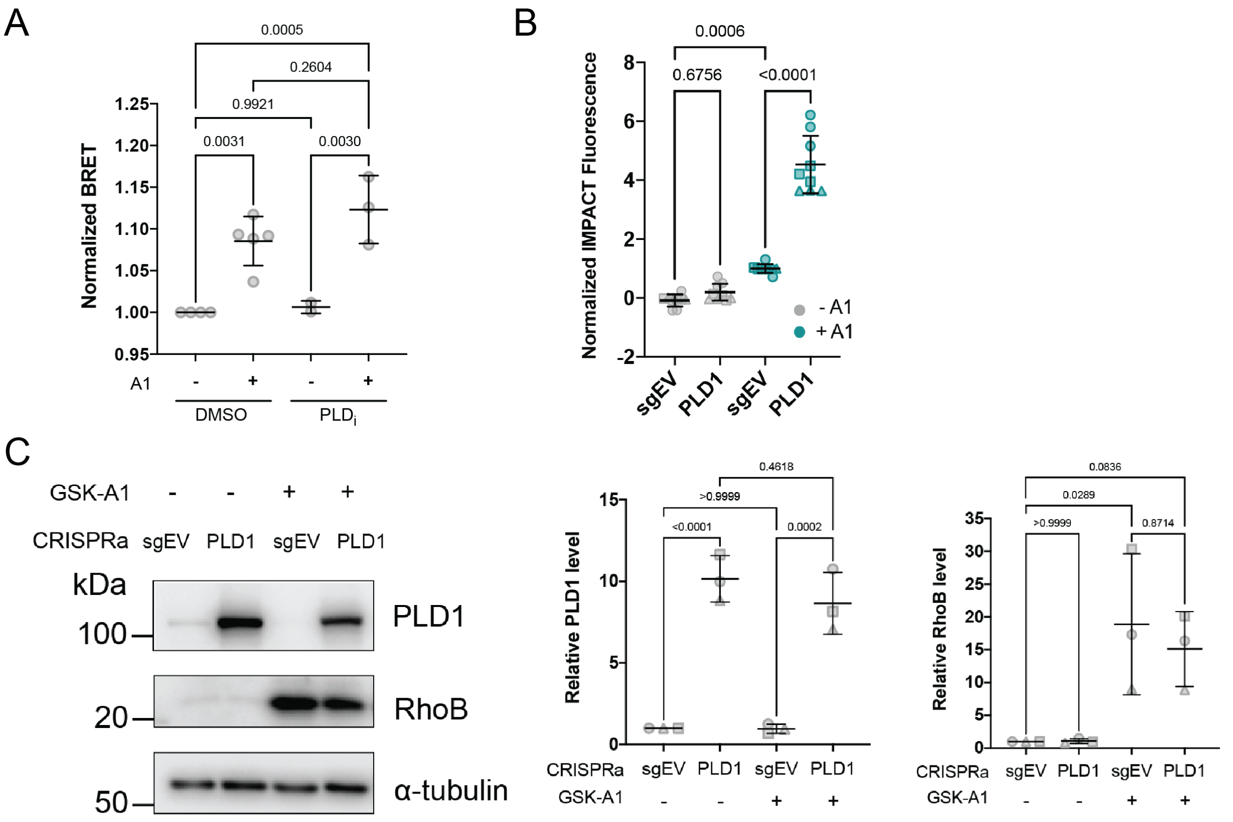


**Figure S2. Effects of GSK-A1 on PM PI(4,5)$P_2$ levels and PLD activity and RhoB levels upon PLD1 CRISPRa.** (A) Live-cell BRET assay analysis of PI(4,5)$P_2$ produced at the PM on HEK293 cells treated with or without GSK-A1 (100 nM, 24 h) or with or without PLD inhibitor FIPI (1 µM) using the $L_{10}$-mVenus-T2A-sLuc-PLCδ$1_{PH}$ PI(4,5)$P_2$ biosensor. n = 5 biological replicates. Statistical significance was determined using one-way ANOVA with Tukey post hoc test, with p values indicated on the plot. (B) Normalized PLD activity by IMPACT on K562 CRISPRa cells with no sgRNA or RHPN2 sgRNAs treated with or without GSK-A1 (100 nM, 18 h). (C) Western blot analysis of K562 cells CRISPRa cells with no sgRNA or RHPN2 sgRNAs treated with or without GSK-A1 (100 nM, 18 h). Quantification performed by normalizing target protein band intensity over Ponceau S intensity. Different shapes on plot correspond to different biological replicates (n = 3). Statistical significance was determined using one-way ANOVA with Tukey post hoc test, with p values indicated on the plot.

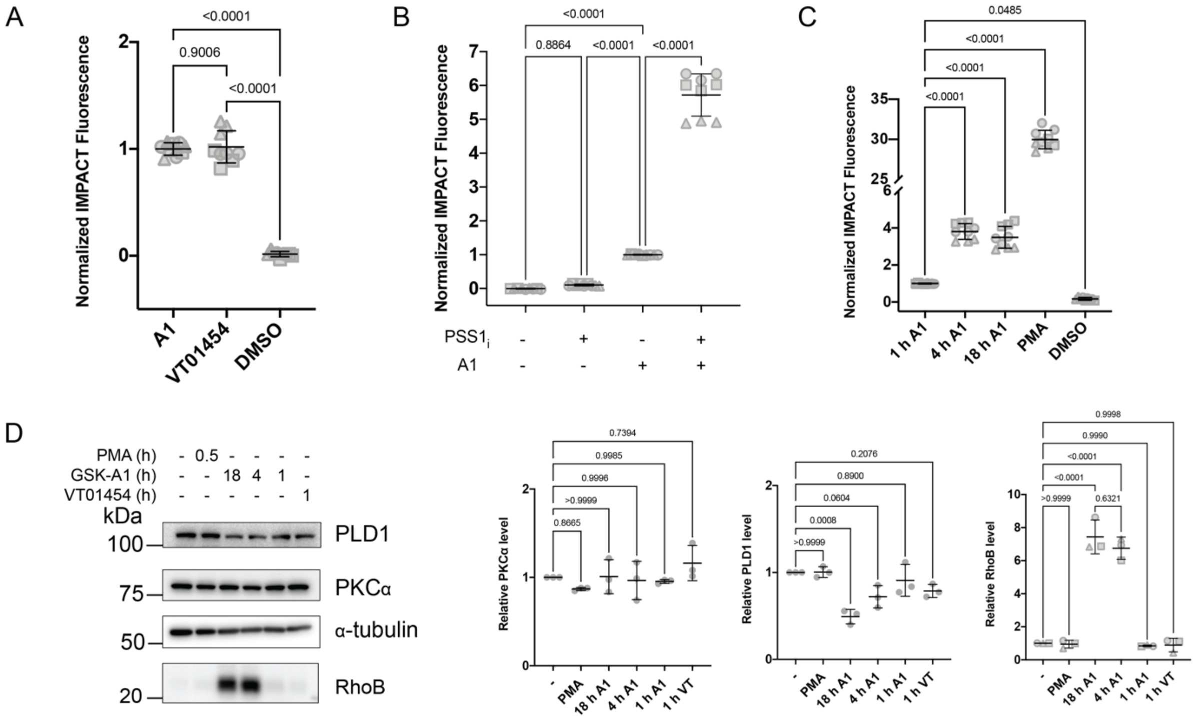


**Figure S3. GSK-A1 effects are recapitulated in U2OS cells.** (A) Normalized PLD activity by IMPACT on HEK293T cells treated with GSK-A1 (100 nM, 1 h), VT01454 (100 nM, 1 h), or DMSO with or without 30 min FIPI pretreatment (750 nM). (B) Normalized PLD activity by IMPACT on U2OS cells pretreated with or without PSS1 inhibitor DS55980254 (1 µM, 24 h) followed by treatment with or without GSK-A1 (100 nM, 4 h). (C) Normalized PLD activity by IMPACT on U2OS with GSK-A1 (100 nM, 0–18 h) or PMA (100 nM, 30 min). (D) Western blot analysis of U2OS cells treated with GSK-A1 (100 nM, 0–18 h), PMA (100 nM, 30 min), or VT01454 (100 nM, 1 h). Quantification performed by normalizing PLD1 or PKCα protein band intensity over Ponceau S intensity. Different shapes on plot correspond to different biological replicates (n = 3). Statistical significance was determined using one-way ANOVA with Tukey post hoc test, with p values indicated on the plot.

| | Gene | ENSEMBL ID | RPF_0min | RPF_60min | RPF_360min | RPF 360/0 ratio |
|---|---|---|---|---|---|---|
| 1 | CCSER2 | ENST00000372088 | 21.66 | 41.13 | 843.41 | 38.94 |
| 2 | HMGB1 | ENST00000341423 | 336.29 | 903.47 | 982.64 | 2.92 |
| 3 | SLC23A2 | ENST00000379333 | 21.66 | 39.76 | 60.81 | 2.81 |
| 4 | TMEM178B | ENST00000565468 | 55.86 | 109.68 | 152.04 | 2.72 |
| 5 | CYP26A1 | ENST00000224356 | 17.10 | 20.56 | 46.41 | 2.71 |
| 6 | HELZ2 | ENST00000467148 | 28.50 | 43.87 | 73.62 | 2.58 |
| 7 | SOX5 | ENST00000537393 | 18.24 | 21.94 | 43.21 | 2.37 |
| 8 | PWWP2B | ENST00000305233 | 10.26 | 12.34 | 24.01 | 2.34 |
| 9 | MPDZ | ENST00000447879 | 49.02 | 53.47 | 112.03 | 2.29 |
| 10 | IDI1 | ENST00000695775 | 114.00 | 141.21 | 252.86 | 2.22 |
| 11 | PARP4 | ENST00000381989 | 10.26 | 19.19 | 22.41 | 2.18 |
| 12 | ACACA | ENST00000614428 | 49.02 | 89.11 | 105.63 | 2.15 |
| 13 | DLX5 | ENST00000648378 | 13.68 | 16.45 | 28.81 | 2.11 |
| 14 | ABCB8 | ENST00000358849 | 18.24 | 35.65 | 38.41 | 2.11 |
| 15 | METTL24 | ENST00000338882 | 66.12 | 75.40 | 137.63 | 2.08 |
| 16 | CLDN23 | ENST00000519106 | 70.68 | 72.66 | 137.63 | 1.95 |
| 17 | NLRP13 | ENST00000342929 | 19.38 | 35.65 | 36.81 | 1.90 |
| 18 | TBC1D12 | ENST00000225235 | 71.82 | 79.52 | 136.03 | 1.89 |
| 19 | OBSL1 | ENST00000404537 | 36.48 | 68.55 | 68.82 | 1.89 |
| 20 | CATSPERB | ENST00000256343 | 323.75 | 581.29 | 590.55 | 1.82 |
| 21 | WDR44 | ENST00000254029 | 66.12 | 104.19 | 118.43 | 1.79 |
| 22 | TSPAN14 | ENST00000372158 | 29.64 | 47.98 | 52.81 | 1.78 |
| 23 | ZBTB26 | ENST00000373656 | 17.10 | 20.56 | 30.41 | 1.78 |
| 24 | IPMK | ENST00000373935 | 21.66 | 34.27 | 38.41 | 1.77 |
| 25 | VPS13D | ENST00000620676 | 18.24 | 26.05 | 32.01 | 1.75 |
| 26 | SCFD2 | ENST00000401642 | 19.38 | 21.94 | 33.61 | 1.73 |
| 27 | **RHOB** | ENST00000272233 | 213.17 | 337.26 | 366.49 | 1.72 |
| 28 | BPTF | ENST00000582467 | 136.80 | 198.79 | 230.46 | 1.68 |
| 29 | SDHA | ENST00000264932 | 17.10 | 23.31 | 28.81 | 1.68 |
| 30 | VPS13B | ENST00000357162 | 34.20 | 53.47 | 57.61 | 1.68 |
| 31 | TRPV4 | ENST00000675670 | 29.64 | 30.16 | 49.61 | 1.67 |
| 32 | GPR153 | ENST00000377893 | 31.92 | 47.98 | 52.81 | 1.65 |
| 33 | RXRA | ENST00000481739 | 33.06 | 37.02 | 54.41 | 1.65 |
| 34 | COASY | ENST00000393818 | 27.36 | 41.13 | 44.81 | 1.64 |
| 35 | CCDC167 | ENST00000373408 | 34.20 | 42.50 | 56.01 | 1.64 |
| 36 | FBXO31 | ENST00000311635 | 42.18 | 46.61 | 68.82 | 1.63 |
| 37 | ETV3 | ENST00000368192 | 14.82 | 20.56 | 24.01 | 1.62 |
| 38 | POR | ENST00000706547 | 269.03 | 381.13 | 433.71 | 1.61 |
| 39 | TFIP11 | ENST00000407690 | 31.92 | 35.65 | 51.21 | 1.60 |
| 40 | ZNF276 | ENST00000443381 | 23.94 | 28.79 | 38.41 | 1.60 |
| 41 | PFAS | ENST00000314666 | 42.18 | 45.24 | 67.22 | 1.59 |
| 42 | PTPN3 | ENST00000374541 | 18.24 | 24.68 | 28.81 | 1.58 |
| 43 | CDCA7L | ENST00000406877 | 125.40 | 168.63 | 196.85 | 1.57 |
| 44 | ANLN | ENST00000265748 | 49.02 | 49.35 | 76.82 | 1.57 |
| 45 | EXOC8 | ENST00000366645 | 69.54 | 71.29 | 108.83 | 1.56 |
| 46 | SGSM2 | ENST00000268989 | 30.78 | 45.24 | 48.01 | 1.56 |

| | | | | | | |
|---|---|---|---|---|---|---|
| 47 | HDAC6 | ENST00000443563 | 20.52 | 30.16 | 32.01 | 1.56 |
| 48 | SPATA2L | ENST00000289805 | 31.92 | 41.13 | 49.61 | 1.55 |
| 49 | SLC39A4 | ENST00000301305 | 28.50 | 28.79 | 43.21 | 1.52 |
| 50 | INTS14 | ENST00000313182 | 85.50 | 89.11 | 128.03 | 1.50 |
| 51 | USP24 | ENST00000294383 | 57.00 | 57.58 | 84.82 | 1.49 |
| 52 | MBD6 | ENST00000355673 | 19.38 | 20.56 | 28.81 | 1.49 |
| 53 | PHLPP2 | ENST00000568954 | 21.66 | 24.68 | 32.01 | 1.48 |
| 54 | FOXD1 | ENST00000615637 | 25.08 | 32.90 | 36.81 | 1.47 |
| 55 | **GADD45B** | ENST00000215631 | 38.76 | 54.84 | 56.01 | 1.45 |
| 56 | WRNIP1 | ENST00000380773 | 103.74 | 135.73 | 148.84 | 1.43 |
| 57 | SLC12A4 | ENST00000316341 | 13.68 | 16.45 | 19.20 | 1.40 |
| 58 | AKAP11 | ENST00000025301 | 30.78 | 37.02 | 43.21 | 1.40 |
| 59 | INF2 | ENST00000392634 | 58.14 | 58.95 | 81.62 | 1.40 |
| 60 | ZCCHC14 | ENST00000671377 | 27.36 | 34.27 | 38.41 | 1.40 |
| 61 | RTTN | ENST00000640769 | 34.20 | 43.87 | 48.01 | 1.40 |
| 62 | GLOD4 | ENST00000301329 | 149.34 | 204.27 | 204.85 | 1.37 |
| 63 | KIAA0556 | ENST00000261588 | 43.32 | 49.35 | 59.21 | 1.37 |
| 64 | CCT7 | ENST00000258091 | 1334.90 | 1416.21 | 1814.85 | 1.36 |
| 65 | HOXB6 | ENST00000484302 | 90.06 | 91.85 | 121.63 | 1.35 |
| 66 | **MELK** | ENST00000298048 | 58.14 | 65.81 | 78.42 | 1.35 |
| 67 | RAPH1 | ENST00000308091 | 57.00 | 64.44 | 76.82 | 1.35 |
| 68 | FUBP3 | ENST00000319725 | 375.05 | 404.44 | 504.12 | 1.34 |
| 69 | **MARK2** | ENST00000502399 | 79.80 | 100.08 | 107.23 | 1.34 |
| 70 | DHX9 | ENST00000367549 | 99.18 | 111.05 | 132.83 | 1.34 |
| 71 | USP39 | ENST00000323701 | 72.96 | 76.77 | 97.62 | 1.34 |
| 72 | PPP1R10 | ENST00000376511 | 47.88 | 49.35 | 64.02 | 1.34 |
| 73 | SLC6A6 | ENST00000622186 | 23.94 | 31.53 | 32.01 | 1.34 |
| 74 | EZR | ENST00000337147 | 70.68 | 82.26 | 94.42 | 1.34 |
| 75 | NDUFA6 | ENST00000498737 | 223.43 | 264.60 | 296.07 | 1.33 |
| 76 | CDK13 | ENST00000181839 | 33.06 | 38.39 | 43.21 | 1.31 |
| 77 | ZNF469 | ENST00000565624 | 63.84 | 72.66 | 83.22 | 1.30 |
| 78 | PRRG2 | ENST00000246794 | 30.78 | 34.27 | 40.01 | 1.30 |
| 79 | ARHGAP20 | ENST00000260283 | 13.68 | 16.45 | 17.60 | 1.29 |
| 80 | TRIM61 | ENST00000508856 | 132.24 | 153.55 | 166.44 | 1.26 |
| 81 | BRCA1 | ENST00000357654 | 120.84 | 131.61 | 152.04 | 1.26 |
| 82 | PDK3 | ENST00000379162 | 42.18 | 52.10 | 52.81 | 1.25 |
| 83 | POLL | ENST00000370162 | 20.52 | 20.56 | 25.61 | 1.25 |
| 84 | VEZT | ENST00000436874 | 55.86 | 57.58 | 68.82 | 1.23 |
| 85 | NID1 | ENST00000264187 | 106.02 | 127.50 | 129.63 | 1.22 |
| 86 | VARS1 | ENST00000375663 | 86.64 | 87.74 | 105.63 | 1.22 |
| 87 | MDN1 | ENST00000369393 | 77.52 | 85.00 | 94.42 | 1.22 |
| 88 | RPLP0 | ENST00000551150 | 1607.36 | 1791.86 | 1955.68 | 1.22 |
| 89 | KCNG2 | ENST00000316249 | 17.10 | 20.56 | 20.81 | 1.22 |
| 90 | ATM | ENST00000452508 | 74.10 | 87.74 | 89.62 | 1.21 |
| 91 | CEP170B | ENST00000414716 | 45.60 | 50.73 | 54.41 | 1.19 |
| 92 | CXXC1 | ENST00000285106 | 71.82 | 72.66 | 84.82 | 1.18 |
| 93 | NRIP1 | ENST00000318948 | 34.20 | 35.65 | 40.01 | 1.17 |
| 94 | MUC19 | ENST00000454784 | 27.36 | 28.79 | 32.01 | 1.17 |

| | | | | | | |
|---|---|---|---|---|---|---|
| 95 | SCAF4 | ENST00000286835 | 117.42 | 117.90 | 136.03 | 1.16 |
| 96 | EIF5B | ENST00000289371 | 45.60 | 50.73 | 52.81 | 1.16 |
| 97 | MYBPC3 | ENST00000545968 | 19.38 | 20.56 | 22.41 | 1.16 |
| 98 | IGSF9B | ENST00000533871 | 12.54 | 13.71 | 14.40 | 1.15 |
| 99 | LTN1 | ENST00000389194 | 25.08 | 28.79 | 28.81 | 1.15 |
| 100 | KCNC4 | ENST00000438661 | 343.13 | 359.19 | 388.90 | 1.13 |
| 101 | RALGAPA2 | ENST00000202677 | 28.50 | 30.16 | 32.01 | 1.12 |
| 102 | MYC | ENST00000524013 | 265.61 | 274.19 | 297.67 | 1.12 |
| 103 | INCENP | ENST00000394818 | 502.73 | 533.31 | 550.54 | 1.10 |
| 104 | FOXE3 | ENST00000335071 | 193.79 | 205.65 | 211.25 | 1.09 |
| 105 | TYK2 | ENST00000525621 | 111.72 | 115.16 | 120.03 | 1.07 |
| 106 | ZNF131 | ENST00000682664 | 47.88 | 49.35 | 51.21 | 1.07 |

**Figure S4. Ribosome profiling results for GSK-A1-treated HEK293T cells.** Ribosome profiling results on HeLa cells treated with GSK-A1 (100 nM, 0–6 h). Gene hits shown are genes with increased RNA levels from 0 to 1 h and 1 h to 6 h over two biological replicates. Genes ranked by ratio of RNA level at 6 h over 0 h of replicate 1. RhoB is shown in red, and MAPK pathway-related genes are colored in blue. Full dataset provided in **Table S2**.

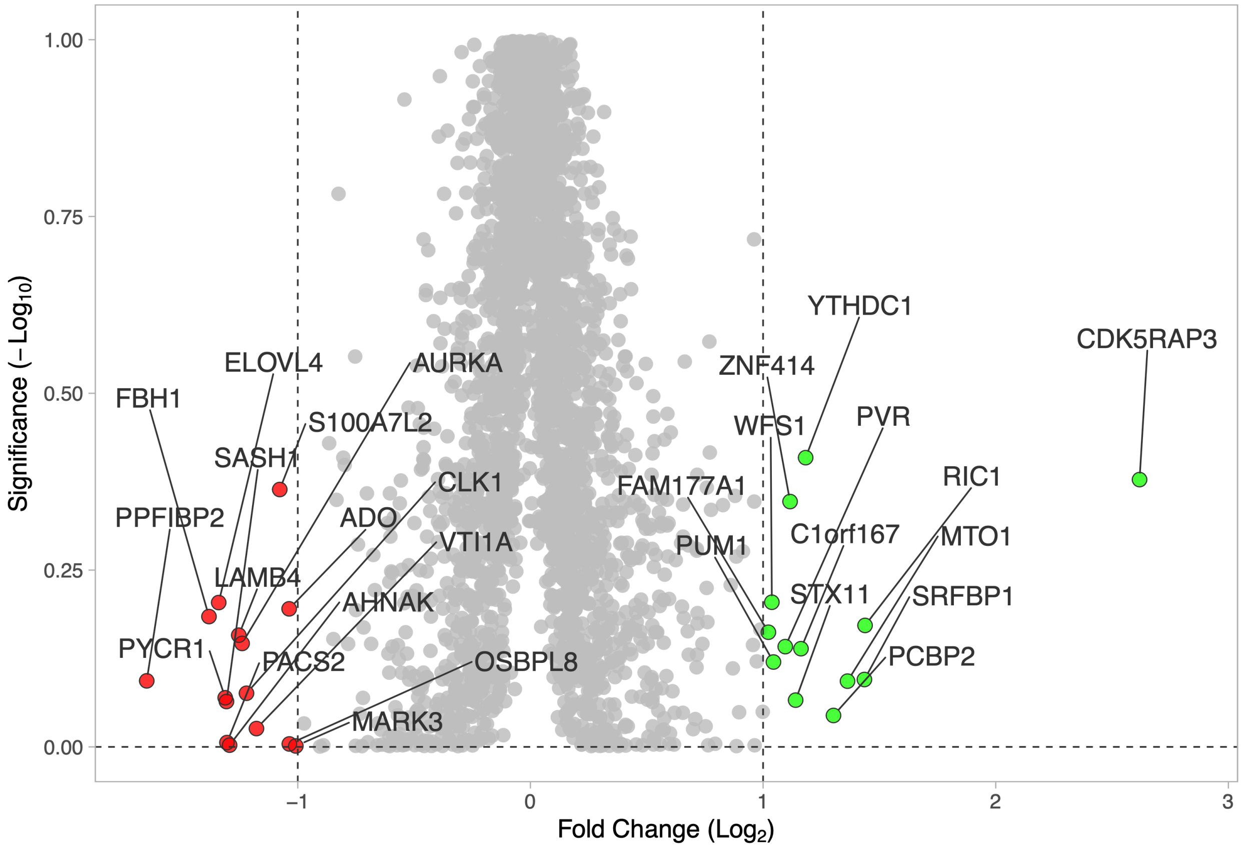


**Figure S5. TurboID proximity labeling proteomics results for GSK-A1-treated HeLa cells.** TurboID proteomics results on HeLa cells stably expressing $Lyn_{10}$-TurboID-V5 treated with or without GSK-A1 (100 nM, 6 h). Green indicates hits enriched and red indicated hits unenriched at the PM after GSK-A1 treatment. Full dataset is provided in **Table S3**.

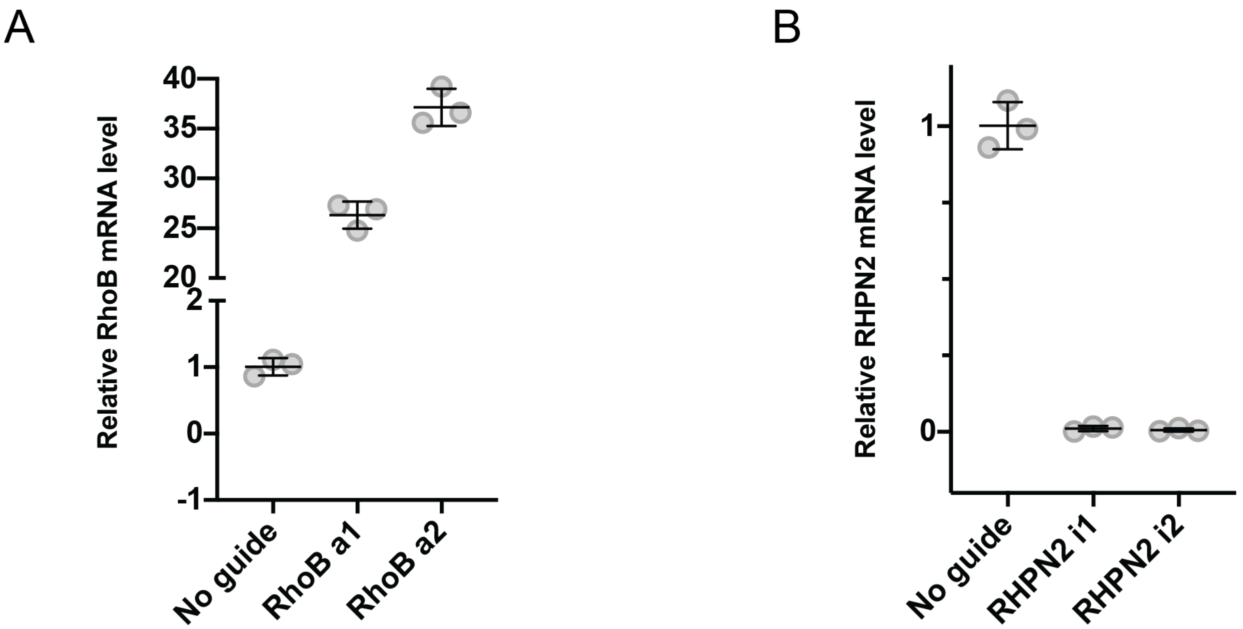


**Figure S6. qPCR validation for CRISPRa/i cell lines.** (A) Relative mRNA levels of K562 CRISPRa-edited RhoB cell lines assessed by qPCR. Values normalized to GAPDH control. (B) Relative mRNA levels of K562 CRISPRi-edited RHPN2 cell lines assessed by qPCR. Values normalized to GAPDH control.

**Table S1. Full dataset from RNA-seq experiments. (.xlsx file)**

**Table S2. Full dataset from ribosome profiling experiments. (.xlsx file)**

**Table S3. Full dataset from proximity labeling experiments. (.xlsx file)**
